# Chemogenomic profiling of anti-leishmanial efficacy and resistance in the related kinetoplastid parasite *Trypanosoma brucei*

**DOI:** 10.1101/605873

**Authors:** Clare F Collett, Carl Kitson, Nicola Baker, Heather B. Steele-Stallard, Marie-Victoire Santrot, Sebastian Hutchinson, David Horn, Sam Alsford

## Abstract

The arsenal of drugs used to treat leishmaniasis, caused by *Leishmania* spp., is limited and beset by toxicity and emergent resistance. Furthermore, our understanding of drug mode-of-action and potential routes to resistance is limited. Forward genetic approaches have revolutionised our understanding of drug mode-of-action in the related kinetoplastid parasite, *Trypanosoma brucei*. Therefore, we screened our genome-scale *T. brucei* RNAi library in the current anti-leishmanial drugs, sodium stibogluconate (antimonial), paromomycin, miltefosine and amphotericin-B. Identification of *T. brucei* orthologues of the known *Leishmania* antimonial and miltefosine plasma membrane transporters effectively validated our approach, while a cohort of 42 novel drug efficacy determinants provides new insights and serves as a resource. Follow-up analyses revealed the antimonial selectivity of the aquaglyceroporin, TbAQP3. A lysosomal major facilitator superfamily transporter contributes to paromomycin/aminoglycoside efficacy. The vesicle-associated membrane protein, TbVAMP7B, and a flippase contribute to amphotericin-B and miltefosine action, and are potential cross-resistance determinants. Finally, multiple phospholipid-transporting flippases, including the *T. brucei* orthologue of the *Leishmania* miltefosine transporter, a putative β-subunit/CDC50 co-factor, and additional membrane-associated hits, affect amphotericin-B efficacy, providing new insights into mechanisms of drug uptake and action. The findings from this orthology-based chemogenomic profiling approach substantially advance our understanding of anti-leishmanial drug action and potential resistance mechanisms, and should facilitate the development of improved therapies, as well as surveillance for drug-resistant parasites.

**Importance:** Leishmaniasis is a devastating disease caused by the *Leishmania* parasites and is endemic to a wide swathe of the tropics and sub-tropics. While there are drugs available for the treatment of leishmaniasis, these suffer from various challenges, including the spread of drug resistance. Our understanding of anti-leishmanial drug action and the modes of drug resistance in *Leishmania* is limited. The development of genetic screening tools in the related parasite, *Trypanosoma brucei*, has revolutionised our understanding of these processes in this parasite. Therefore, we applied these tools to the anti-leishmanial drugs, identifying *T. brucei* orthologues of known *Leishmania* proteins that drive drug uptake, as well as a panel of novel proteins not previously associated with anti-leishmanial drug action. Our findings substantially advance our understanding of anti-leishmanial mode-of-action and provide a valuable starting point for further research.

## Introduction

The kinetoplastid parasites, *Leishmania* species, *Trypanosoma brucei* subspecies and *T. cruzi* are respectively endemic throughout much of the tropics and sub-tropics, sub-Saharan Africa and Latin America. They are responsible for various forms of leishmaniasis (*Leishmania* spp.) (1), human African trypanosomiasis (HAT; *T. b. gambiense* and *T. b. rhodesiense*) and the livestock disease, nagana (*T. b. brucei* and related African trypanosomes) (2), and Chagas’ disease (*T. cruzi*) (3). Collectively, these parasites cause a huge burden of disease amongst predominantly poor populations in affected regions. Leishmaniasis is caused by a range of *Leishmania* species, leading to cutaneous and visceral forms of the disease, of which there are 0.7-1.3 and 0.2-0.4 million cases annually (4). While cutaneous leishmaniasis can be self-limiting, infections with *L. braziliensis* (and other members of the *Viannia* sub-genus) can develop into mucocutaneous leishmaniasis, a profoundly disfiguring form of the disease (4). Visceral leishmaniasis, also known as kala-azar, is typically fatal if untreated.

There are four current anti-leishmanial drugs: sodium stibogluconate (SSG), paromomycin, miltefosine and amphotericin-B, which are unsatisfactory due to toxicity, emerging drug resistance, complex administration protocols and variable efficacy depending on the disease type or infecting *Leishmania* species (5). With the exception of miltefosine (in use against leishmaniasis since 2002), the current anti-leishmanial drugs have been in use for many decades. Until recently, efforts have focused on the development of more effective drug delivery regimens and combination therapies, with the aim of reducing dosages (and therefore side effects) and combating the emergence of resistance. The rise of antimonial resistant *L. donovani* on the Indian sub-continent now precludes the use of sodium stibogluconate (SSG) (6), while miltefosine resistant *L. donovani* has been confirmed in the clinic (7). Consequently, the World Health Organisation recommends various combination therapies, depending on the *Leishmania* species and geographical region (8). However, it is relatively easy to generate *Leishmania* resistant to combination therapies in the laboratory (9, 10). More recently, new drugs have entered the clinical development pipeline. However, the most advanced of these, fexinidazole, which recently passed phase 2/3 clinical trials against HAT (11), and has anti-leishmanial activity *in vitro* (12), lacks efficacy *in vivo* (13).

Given the ease with which *Leishmania* parasites become resistant to the available drugs, it is critically important to understand how this resistance might develop. Identification of the genetic changes underlying drug resistance will enable the development of molecular diagnostics to inform treatment choice (14). *Leishmania* genome and transcriptome analyses have identified large numbers of candidate genes (15, 16) but relatively few have been directly linked to drug-action. While some drugs can freely move across membranes, many are taken up *via* specific surface receptors and transporters. For example, miltefosine uptake is dependent on a *Leishmania* amino phospholipid-transporting (P4)-type ATPase (or flippase) and its β-subunit/CDC50 co-factor, Ros3 (17, 18), while the Sb(III) form of SSG is taken up *via* an aquaglyceroporin, AQP1 (19). There is also evidence that the ABC transporter, MRPA, influences SSG uptake and sequestration (20), and several other proteins have been implicated in SSG efficacy (reviewed in (14)). In addition, the generation of drug resistant *Leishmania* in the laboratory and various ‘omics analyses have provided insights into anti-leishmanial drug action and resistance mechanisms. Proteomic analyses of paromomycin resistant *L. donovani* revealed a complex picture, with a range of proteins upregulated, including several involved in translation regulation, vesicular trafficking and glycolysis (21). A similar analysis of amphotericin-B resistant *L. infantum* highlighted the differential expression of metabolic enzymes and the upregulation of proteins involved in protection against reactive oxygen species (22). Metabolomic analyses suggested that oxidative defence also contributes to SSG/amphotericin-B and SSG/paromomycin resistance in *L. donovani* (23).

The studies described above highlight the phenotypic consequences of changes in drug sensitivity, but not necessarily the genetic changes responsible. Forward genetic approaches can identify genes that contribute to drug action and resistance. For example, genome-scale RNAi library screening, coupled with RNA interference target sequencing (RIT-seq), has revolutionised our understanding of anti-HAT drug action and resistance (24, 25). In addition, the Cos-seq approach has enabled gain-of-function screening in *Leishmania* (26), leading to target validation for *N*-myristoyltransferase (27) and the identification of a panel of putative antimony and miltefosine resistance genes (28). While undoubtedly a powerful technique, Cos-seq is unable to identify drug uptake or activation mechanisms, which can be characterised by loss-of-function approaches, such as RIT-seq. However, due to the absence of the RNAi machinery in most *Leishmania* species (with the notable exception of *L. braziliensis* (29)), this loss-of-function approach is not possible in these parasites. Although *T. brucei* and *Leishmania* have distinct life cycles, they are phylogenetically related kinetoplastid parasites that exhibit a high degree of biochemical and genetic similarity (30). Indeed, the majority of orthologous genes are syntenic, indicating little change in gene-order since divergence from a common ancestor. Perhaps not surprisingly then, several ‘dual-purpose’ drugs display activity against both parasites, including pentamidine (5), fexinidazole (11, 12) and the proteasome inhibitor, GNF6702 (31). *T. brucei* is also susceptible to *in vitro* killing by the four current anti-leishmanial drugs. Therefore, we hypothesised that *T. brucei* RNAi library selection in the anti-leishmanial drugs would enable identification of candidate drug efficacy determinants with orthologues in *Leishmania*.

Here, we describe RIT-seq library screening using each of the current anti-leishmanial drugs. We identified 44 high confidence putative drug efficacy determinants, including the *T. brucei* orthologues of the *Leishmania* SSG and miltefosine transporters. Among many previously unknown drug efficacy determinants, we found that the vesicle-associated membrane protein, TbVAMP7B, contributes to miltefosine and amphotericin-B efficacy, and highlight a role for a cohort of amino phospholipid-transporting P4-ATPases (or ‘flippases’) in driving amphotericin-B efficacy. This collection of validated and putative anti-leishmanial drug efficacy determinants provides new insight into mode-of-action and potential resistance mechanisms, and represents an important resource to guide future study.

## Results

### Orthology-based chemogenomic profiles for anti-leishmanial drugs

The four current anti-leishmanial drugs, sodium stibogluconate (SSG), paromomycin, miltefosine and amphotericin-B, have *in vitro* EC_50_ values against *T. brucei* of 1.8 µg.ml^−1^, 17 µM, 30 µM and 260 nM, respectively (Fig. 1A). The equivalent values versus intracellular *L. donovani* amastigotes in mouse peritoneal macrophages are approximately an order of magnitude higher (SSG, paromomycin) or lower (miltefosine, amphotericin-B) (32). To identify factors whose loss renders *T. brucei* less sensitive to each anti-leishmanial drug, a bloodstream-form (BSF) *T. brucei* RNAi library was induced for 24 hours then each drug added at 1-3X EC_50_; selection and induction were maintained thereafter (Fig. 1B). After selection for approximately 10 days, populations with reduced drug sensitivity emerged and grew consistently under continued selection (Fig. 1C).

**Figure 1.**
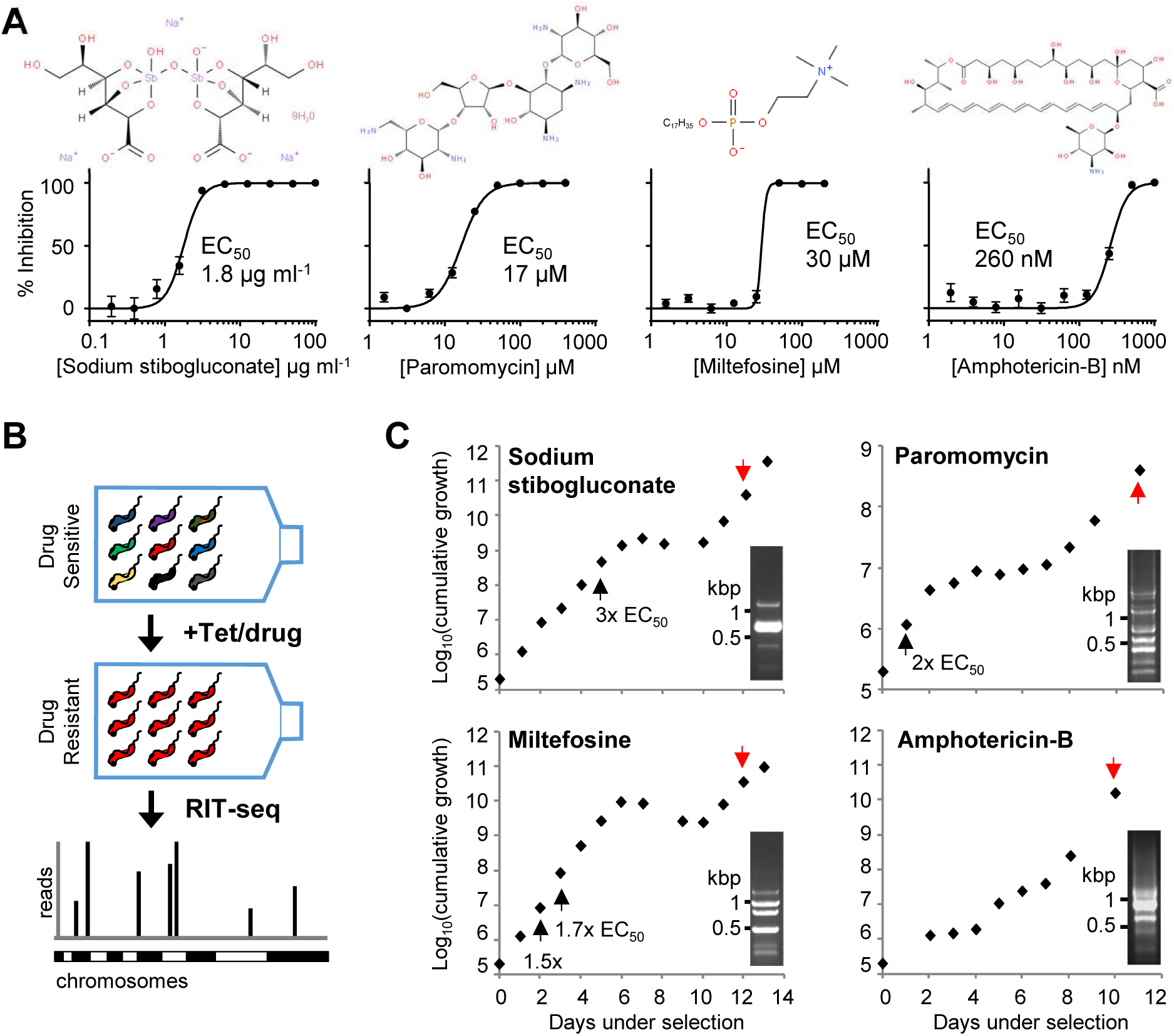
Anti-leishmanial drug selection of a genome-scale *T. brucei* RNAi library. A) Representative EC_50_ charts showing the susceptibility of *T. brucei* to the anti-leishmanial drugs. Individual EC_50_ assays were carried out in quadruplicate; error bars represent standard deviation. Insets: structures of the anti-leishmanial drugs (www.Chemspider.com). B) Schematic showing bloodstream-form *T. brucei* RNAi library selection and RNAi fragment identification by RIT-seq. C) Growth during anti-leishmanial drug selection of the BSF *T. brucei* RNAi library; selection was initiated in 1.5X EC_50_, except for miltefosine (1.0X EC_50_), and adjusted as indicated (black arrows); induction in 1 µg.ml^−1^ tetracycline was maintained throughout. Genomic DNA prepared at the indicated times (red arrows). Insets: RNAi library-specific PCR.

Following robust growth for at least two days, genomic DNA was isolated from the drug-selected populations and subjected to RNAi construct-specific PCR, generating distinct banding patterns for each (Fig. 1C). We sequenced the amplified RNAi target fragment populations from the selected RNAi libraries on an Illumina HiSeq platform (Table S1). For each selected RNAi library, we mapped more than three million individual sequence reads, representing anti-leishmanial enriched RNAi target fragments, to the TREU927 *T. brucei* reference genome (33) using our established RIT-seq methodology (34) (Fig. 1B). The presence of the RNAi construct-specific barcode identified ‘high confidence’ hits, i.e. those represented by more than 99 barcoded reads/kilobase/predicted transcript (open reading frames plus predicted untranslated regions, as annotated in the TREU927 reference genome available at www.tritrypdb.org), and recovery of at least two independent RNAi target fragments (Fig. 2; Fig. S1; Table S1).

**Figure 2.**
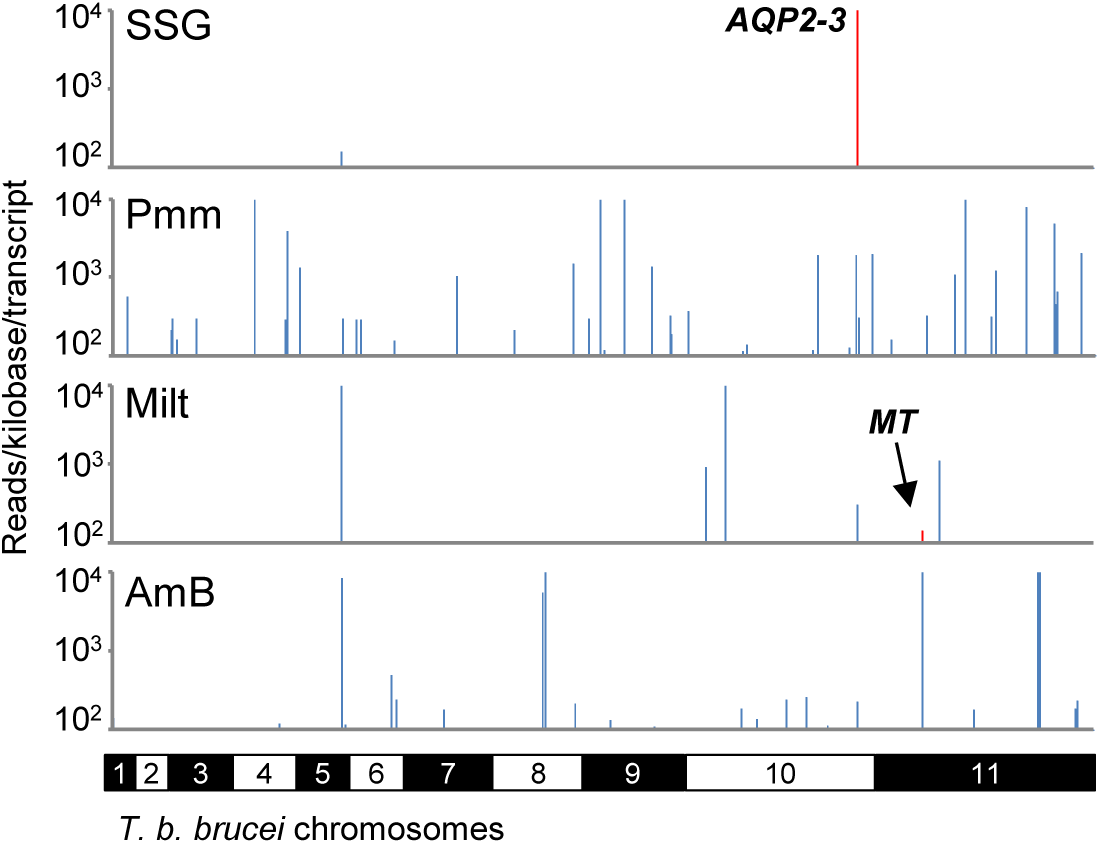
Genome-scale maps showing hits in each screen. Illumina sequencing of the amplified RNAi target fragments identifies *T. brucei* orthologues of known *Leishmania* drug transporters and novel putative drug efficacy determinants. RNAi fragments amplified from each selective screen were mapped against the TREU927 *T. brucei* reference genome. Red bars correspond to *T. brucei* orthologues of known *Leishmania* drug transporters: *AQP2-3*, aquaglyceroporin-2-3 locus, Tb927.10.14160-70; *MT*, miltefosine transporter orthologue, Tb927.11.3350. The y-axes are truncated to 10^4^ reads/kilobase/transcript. SSG, sodium stibogluconate; Pmm, paromomycin; Milt, miltefosine; AmB, amphotericin-B.

Importantly, we identified *T. brucei* orthologues of two known *Leishmania* determinants of anti-leishmanial drug efficacy. RNAi target fragments that mapped to the *TbAQP2-3* locus (Tb927.10.14160-70), which encodes two aquaglyceroporins, dominated the SSG-selected RNAi library; *L. donovani* AQP1 (LdBPK_310030.1) is a key mediator of SSG uptake (19). Another significant hit identified following miltefosine selection was a putative flippase (Tb927.11.3350); the corresponding coding sequence is syntenic with the *L. donovani* miltefosine transporter (LdBPK_131590.1) (17). The identification of *T. brucei* orthologues of these known anti-leishmanial efficacy determinants highlights the power of this chemogenomic profiling approach in the identification of mechanisms of action and resistance that are also relevant to *Leishmania* parasites. In addition to these hits, our RIT-seq analyses yielded a further 42 high confidence hits (Fig. 2; Fig. S1; Table S1).

### TbAQP3, an orthologue of *Leishmania* AQP1, is linked to antimonial action

Aquaglyceroporin defects in *T. brucei* and in *Leishmania* have been linked to arsenical and antimonial resistance (see above), but specific relationships among drugs and AQPs have not been fully elucidated. For example, TbAQP2 is responsible for pentamidine and melarsoprol uptake (35), possibly *via* receptor-mediated endocytosis in the former case (36), and mutations that disrupt *TbAQP2* are responsible for melarsoprol resistance in patients (37). *L. donovani* AQP1 has also been linked to antimonial resistance in patients (38). Notably, TbAQP3 and *Leishmania* AQP1 have the same set of selectivity filter residues (NPA/NPA/WGYR), while TbAQP2 has a divergent set (NSA/NPS/IVLL) (39). Therefore, we investigated the specificity of the interaction between SSG and TbAQP2/TbAQP3, the major hits in the SSG screen.

Sequence mapping of the RNAi target fragments following SSG selection revealed that approximately 71% and 29% of mapped reads containing the RNAi construct-specific barcode corresponded to *TbAQP2* (Tb927.10.14170) and *TbAQP3* (Tb927.10.14160), respectively (Fig. 3A); only 0.08% of reads mapped elsewhere in the genome. These data are consistent with the idea that both aquaglyceroporins contribute to SSG action. However, the*TbAQP2* and *TbAQP3* coding sequences are 82.3% identical, thus while an RNAi fragment may unambiguously map to *TbAQP2*, it may be sufficiently similar to *TbAQP3* to elicit its depletion. Therefore, we tested the relative contribution of the encoded aquaglyceroporins to SSG action against *T. brucei* using *aqp2-3* null and re-expression cell lines (35).

**Figure 3.**
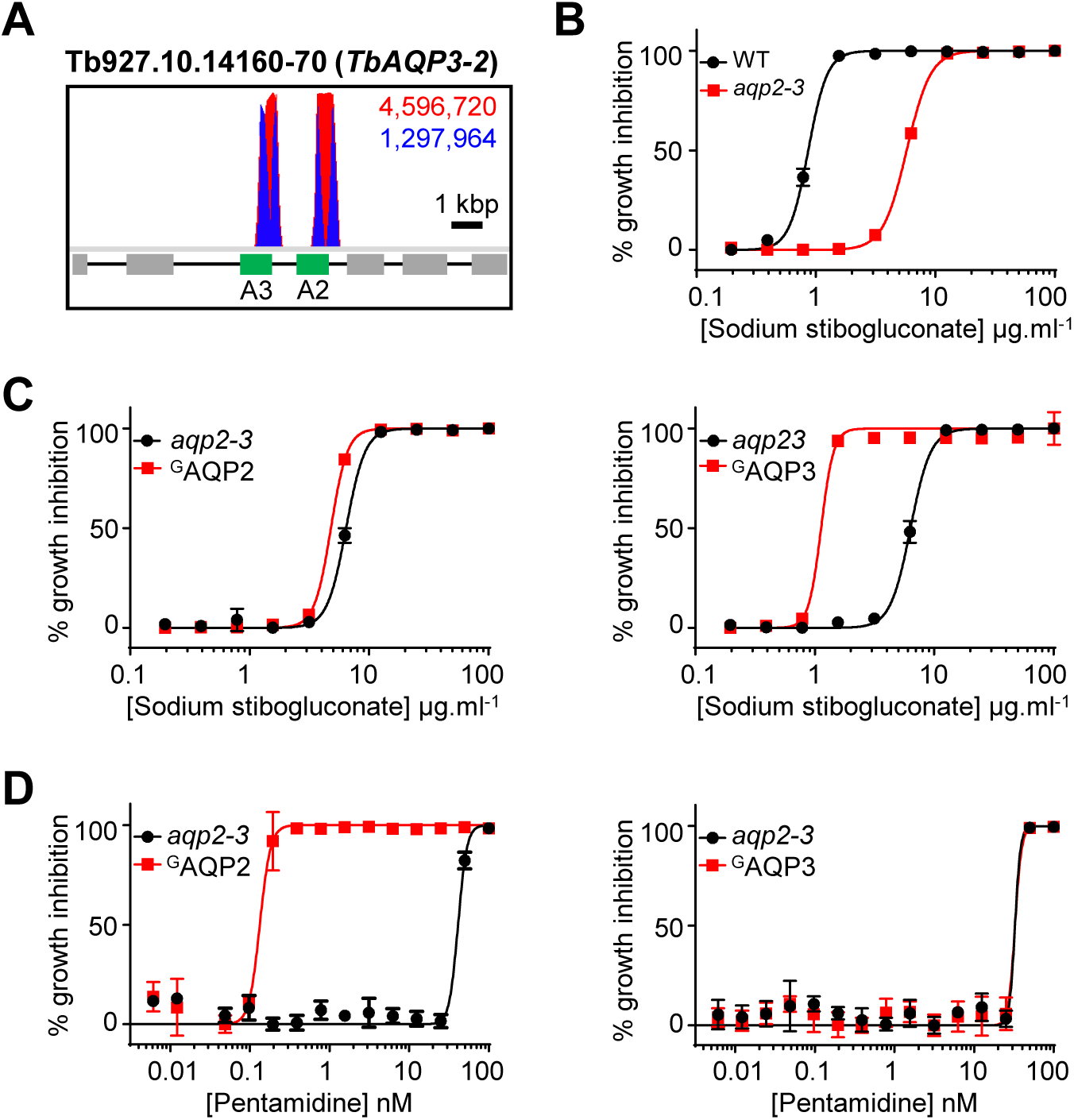
TbAQP3, a *T. brucei* orthologue of *Leishmania* AQP1, is selective for sodium stibogluconate. A) Total (red) and RNAi construct-specific 14mer-containing (blue) reads mapping to the *TbAQP2-3* locus, Tb927.10.14160-70. Targeted open reading frames highlighted in green; flanking open reading frames coloured grey. B) Sodium stibogluconate EC_50_ assay following deletion of the *T. brucei AQP2-3* locus (*aqp2-3*). C) Sodium stibogluconate and D) pentamidine EC_50_ assays following expression of ^GFP^AQP2 (left panels) and ^GFP^AQP3 (right panels) in *aqp2-3* null *T. brucei*. Individual EC_50_ assays were carried out in quadruplicate. Error bars represent standard deviation. WT, *T. brucei* wild type for the *AQP2-3* locus.

Deletion of the *TbAQP2-3* locus led to a 6.7-fold increase in SSG EC_50_ (Fig. 3B), consistent with the output from the screen. Inducible expression of ^GFP^TbAQP2 in the null cell line had little effect on *T. brucei* SSG sensitivity (Fig. 3C, left-hand panel); however, ^GFP^TbAQP3 expression reduced the SSG EC_50_ 5.5-fold (Fig. 3C, right-hand panel). In contrast, and as shown previously (35), ^GFP^AQP2 expression complemented the pentamidine resistance of *aqp2-3* null *T. brucei* (Fig. 3D, left-hand panel), while ^GFP^AQP3 expression had no effect on pentamidine sensitivity (Fig. 3C, right-hand panel). Therefore, SSG sensitivity and resistance is specifically determined by TbAQP3 expression. This indicates that the NPA/NPA/WGYR selectivity filter, present in both TbAQP3 (39) and *Leishmania* AQP1, may be selective for antimonial uptake.

### *T. brucei* lysosomal MFST influences aminoglycoside action

Selection of the BSF *T. brucei* RNAi library with the anti-leishmanial aminoglycoside, paromomycin, identified 50 hits, of which 28 fulfilled our high stringency criteria (Table S1). Twenty-one of the high confidence hits were functionally annotated, and included several associated with transport and nucleic acid processing. The top three hits with functional annotations were *Tb927.9.6360-80* (major facilitator superfamily transporters, MFST), *Tb927.11.6680* (amino acid transporter, AAT15) and *Tb927.11.14190* (Tudor domain-containing Staphylococcal nuclease, TSN) (40), targeted by approximately 84%, 1.7% and 0.9% of the mapped reads, respectively (Fig. 4A, Fig. S2 and Table S1). However, while parasites able to deplete AAT15 and TSN persisted in the population over the 12 days of selection in paromomycin, we were unable to detect a significant advantage versus wild type *T. brucei* during the course of a standard 72-hour EC_50_ assay (Fig. S1). Therefore, we focussed our attention on the MFST genes.

**Figure 4.**
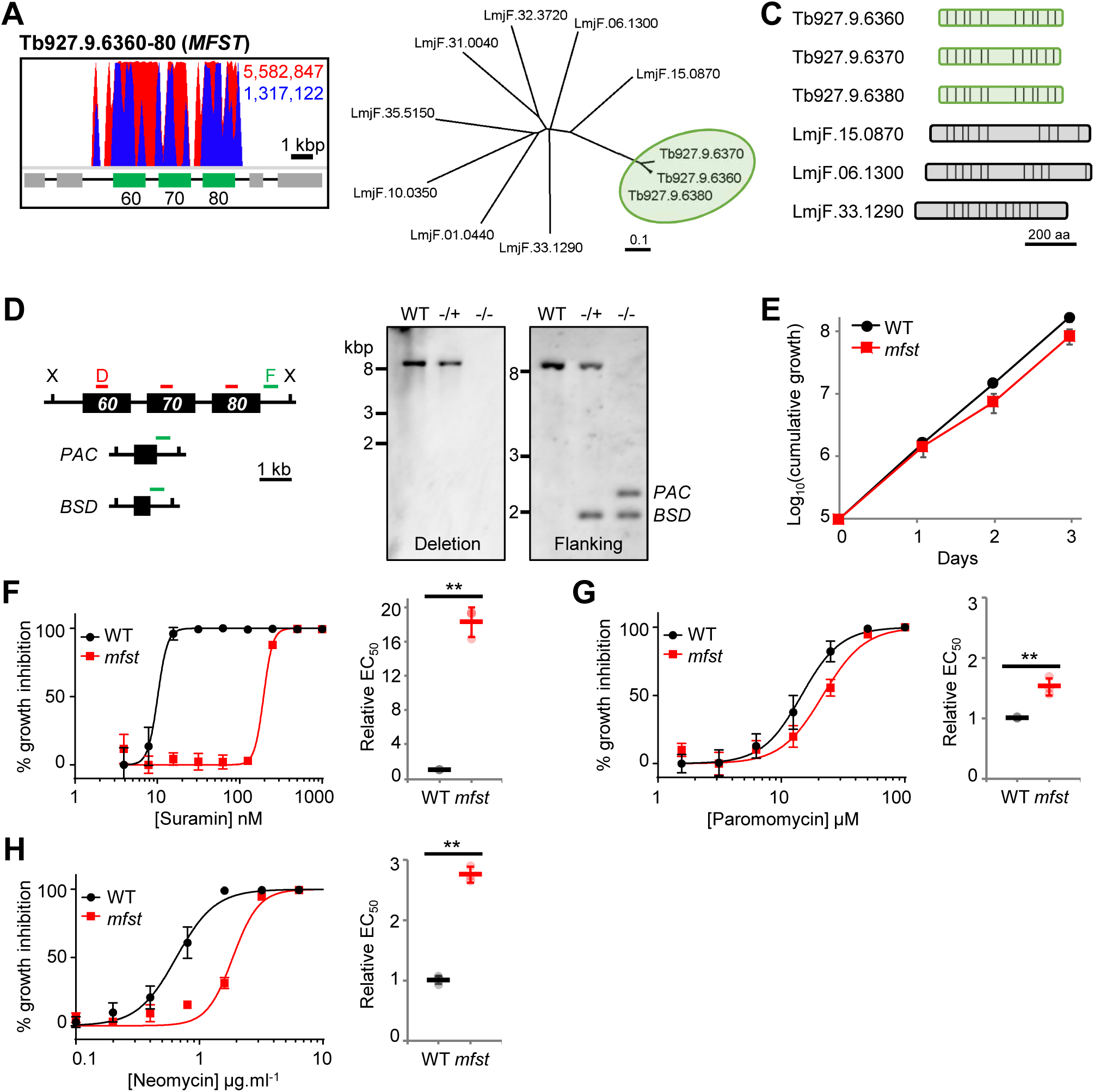
The *T. brucei* lysosomal major facilitator superfamily protein influences the efficacy of aminoglycoside drugs. A) Total (red) and RNAi construct-specific 14mer-containing (blue) reads mapping to the *MFST* locus, *Tb927.9.6360-80*. Targeted open reading frames highlighted in green; flanking open reading frames coloured grey. B) Unrooted neighbour joining tree comparing representative *Leishmania* MFST proteins with Tb927.9.6360-80 (highlighted in green; see Fig. S3 for extended tree). C) Predicted *trans*-membrane organisation of the Tb927.9.6360-80 proteins and the selected *Leishmania* proteins (TM domains, vertical bars). D) *MFST* locus deletion strategy and Southern hybridisation confirming generation of heterozygous (-/+) and homozygous (-/-) *MFST* locus null *T. brucei*. X, *Xho*I; D, deletion probe; F, flanking probe; *PAC*, puromycin acetyltransferase; *BSD*, blasticidin S-deaminase; WT, wild type. E) Growth of WT and *MFST* locus null (*mfst*) *T. brucei* in culture. F-H) Representative EC_50_ assays comparing the sensitivity of WT and *mfst T. brucei* to F) suramin, G) paromomycin and H) neomycin. Inset charts summarise EC_50_ data from three independent biological replicates. Individual growth (E) and EC_50_ (F-H) assays were carried out in triplicate and quadruplicate, respectively. Error bars represent standard deviation. *P*-values derived from Student’s *t*-test (** *P*<0.01).

The genes at the *Tb927.9.6360-80* locus share at least 92% sequence identity and encode for three putative MFSTs, a ubiquitous family of proteins responsible for membrane transit of a wide range of solutes including drugs (41). Comparison with the sequences annotated ‘MFS’ or ‘major facilitator superfamily transporter’ in the *L. major* reference genome confirmed that the syntenic coding sequence, *LmjF.15.0870*, is most closely related to *Tb927.9.6360-80* (Fig. 4B; Fig. S3). The *Leishmania* and *T. brucei* proteins share a similar *trans*-membrane domain organisation and the cytoplasmic loop between TM6 and TM7, which is characteristic of MFST proteins (Fig. 4C) (42).

We previously identified the *Tb927.9.6360-80* locus as a key contributor to suramin efficacy against *T. brucei*, with RNAi depletion of the three transcripts leading to a ten-fold reduction in parasite sensitivity to suramin; localisation studies also indicated that at least one of these transporters is lysosomal (24). Deletion of the whole locus (Fig. 4D) revealed that the three encoded proteins are collectively dispensable in cultured BSF *T. brucei* (Fig. 4E), and enabled us to confirm that not only do these proteins influence suramin efficacy (Fig. 4F), but also that of paromomycin (Fig. 4G) and the related aminoglycoside, neomycin (Fig. 4H). While loss of these MFST proteins dramatically reduces suramin efficacy, the effect on paromomycin and neomycin sensitivity is less pronounced (1.5 and 2.8-fold EC_50_ increase, respectively), though significant. Our mutant BSF *T. brucei* also exhibited better tolerance than wild type parasites to the aminoglycosides at concentrations equivalent to greater than EC_99_ during the first 24 hours of exposure (Fig. S4).

### TbVAMP7B, a cross-efficacy determinant for amphotericin-B and miltefosine

To identify anti-leishmanial cross-efficacy determinants, we next used pairwise comparisons of RNAi library screen outputs (Fig. 5). We first identified a small cohort of hits represented by at least two RNAi target fragments and >99 reads/kilobase/transcript in more than one screen. This group included the *AQP2-3* locus, represented by at least 100 reads in all four screens. We did not explore this locus further since the read-count was at least three orders of magnitude lower in each screen relative to the SSG screen, and leishmanial AQPs have not been implicated in resistance to the other drugs (see above). Two other loci fulfilled our stringency criteria, and both were enriched following amphotericin-B and miltefosine selection: *Tb927.5.3550-70* and *Tb927.11.3350* (Table S1); further analysis of the former hit is considered in this section, while the contribution of Tb927.11.3350 to drug action is addressed subsequently.

**Figure 5.**
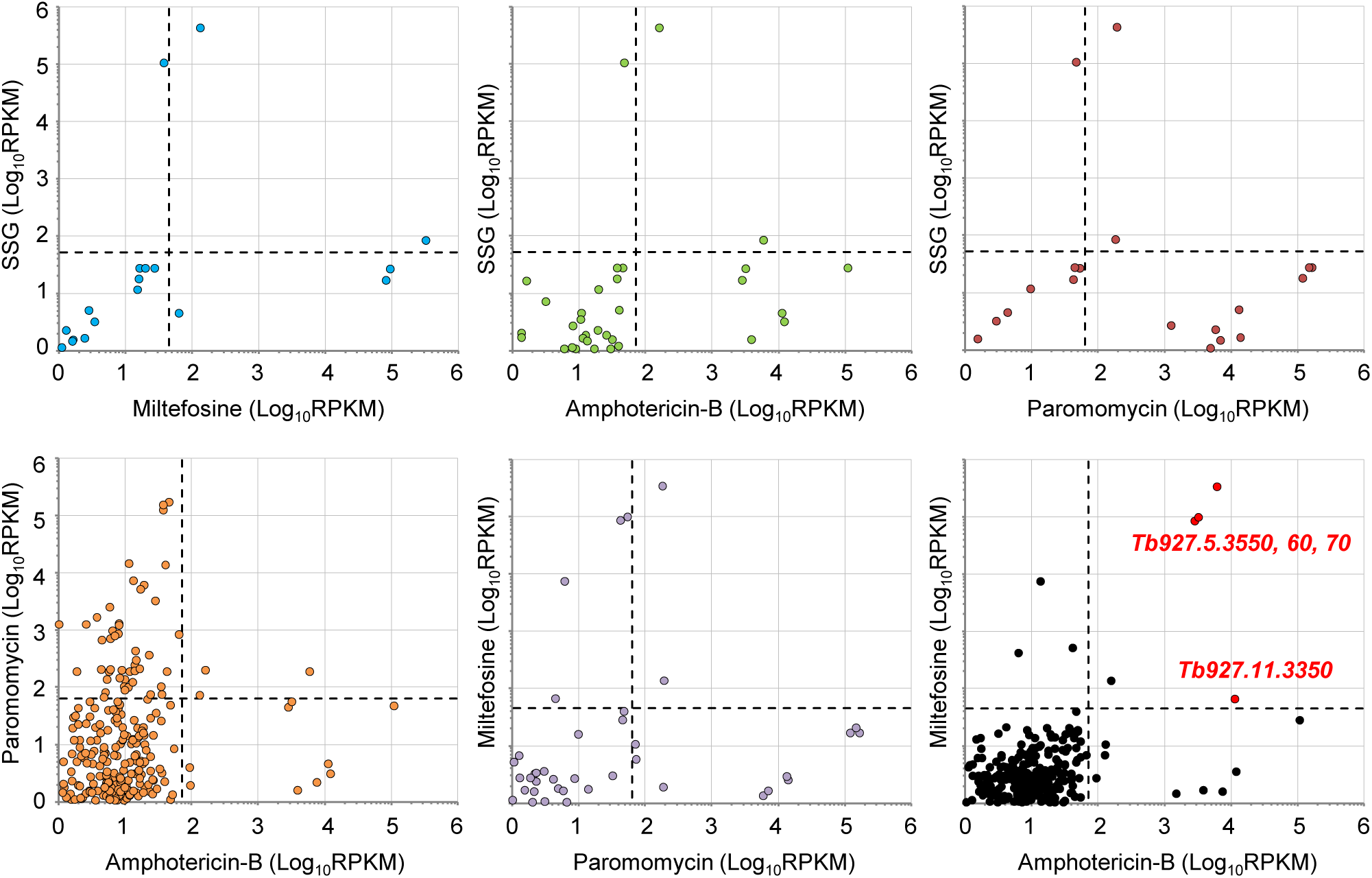
Pairwise comparisons identify putative amphotericin-B/miltefosine cross-efficacy loci. Pairwise comparisons of the sequenced outputs from the four selective screens. Data converted to <u>r</u>eads <u>p</u>er <u>k</u>ilobase per <u>m</u>illion mapped reads (RPKM) to control for minor inter-library variations in read depth. Dashed lines represent stringent 100-read cut offs for each selected RNAi library converted to RPKM. High confidence cross-efficacy determinants highlighted in red in the top right quadrant following comparison of the miltefosine and amphotericin-B selected RNAi libraries.

RIT-seq analysis revealed that 2.2% and 97% of mapped reads identified *Tb927.5.3550-70* in the amphotericin-B and miltefosine screens, respectively (Fig. 6A). This locus encodes for a thioredoxin-like protein (*Tb927.5.3550*), a vesicle-associated membrane protein, TbVAMP7B (*Tb927.5.3560*) (43), and a hypothetical protein (*Tb927.5.3570*). Analysis of the RNAi target fragments mapping to *Tb927.5.3550-70* revealed that few uniquely targeted the *TbVAMP7B* coding sequence (Fig. 6A). Instead, the RNAi target fragments that mapped to the flanking genes overlapped either the *TbVAMP7B* coding sequence (*3550* RNAi target fragments) or 3’-untranslated region (*3570* RNAi target fragments). This pattern is consistent with poor tolerance of TbVAMP7B depletion. Our previous high-throughput phenotypic analysis indicated that TbVAMP7B RNAi knockdown is associated with a significant loss of fitness, while depletion of the flanking transcripts had a less dramatic effect (Table S1) (44). Taken together, these data suggested that TbVAMP7B is an amphotericin-B/miltefosine cross-efficacy determinant, while the identification of the flanking genes was due to bystander effects.

**Figure 6.**
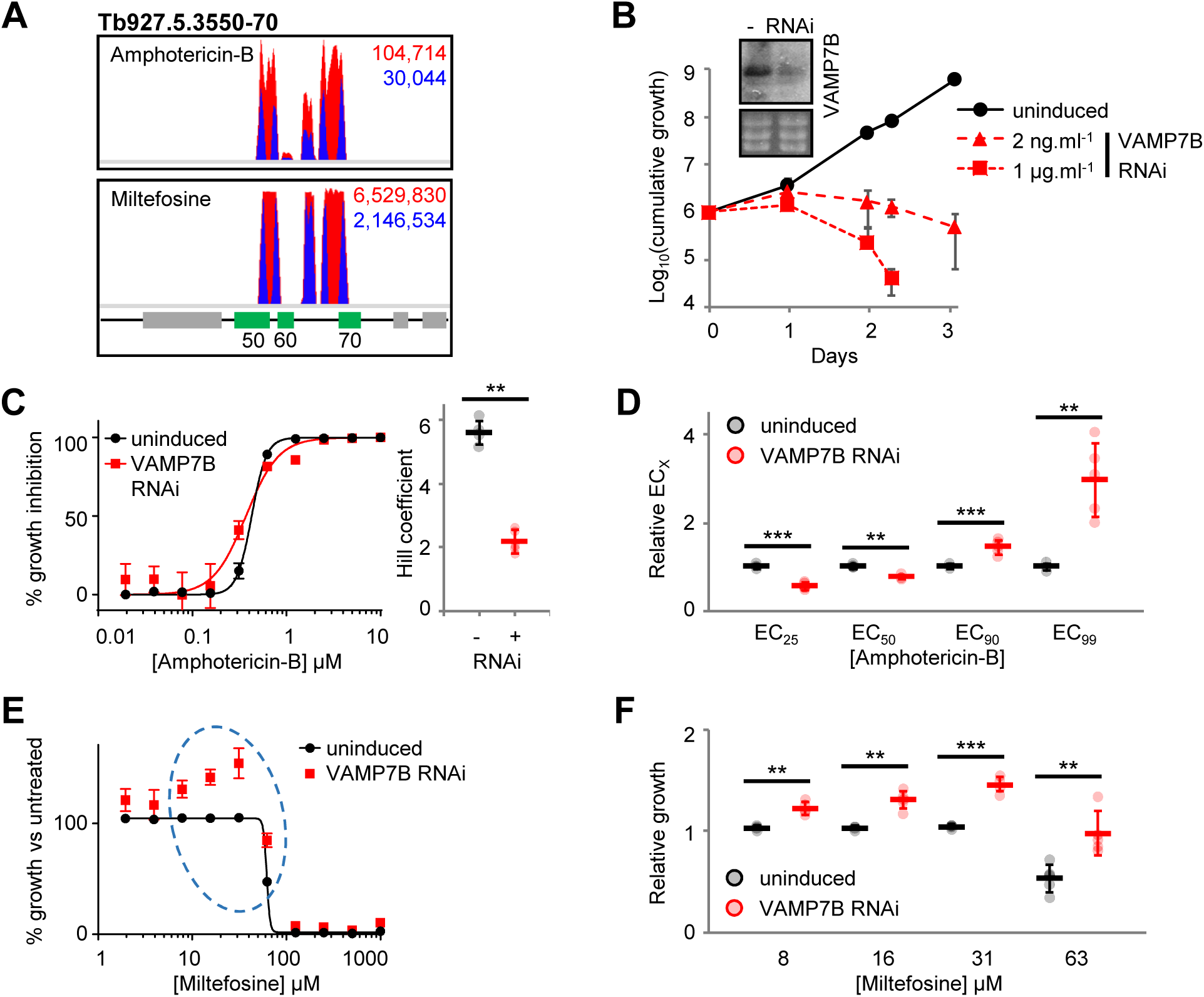
*T. brucei* VAMP7B, Tb927.5.3560, and the action of amphotericin-B and miltefosine. A) Total (red) and RNAi construct-specific 14mer-containing (blue) reads mapping to *Tb927.5.3550-70* following amphotericin-B and miltefosine selection. Targeted open reading frames highlighted in green; flanking open reading frames coloured grey. B) *T. brucei* population growth following TbVAMP7B (Tb927.5.3560) RNAi knockdown. Inset: confirmation of RNAi knockdown by northern blot following 24-hour induction in 1 µg.ml^−1^ tetracycline; ethidium bromide stained gel shown as a loading control. C) Representative 30-hour amphotericin-B EC_50_ assay following TbVAMP7B RNAi knockdown induced in 2 ng.ml^−1^ tetracycline. Inset chart summarises Hill coefficient data for five biological replicates. D) The effect of TbVAMP7B RNAi knockdown on EC_X_ for five biological replicates; data derived for each replicate from EC_50_ values and Hill coefficients presented in (C). E) Representative 30-hour miltefosine EC_50_ assay following TbVAMP7B RNAi knockdown induced in 2 ng.ml^−1^ tetracycline; data plotted to show population growth relative to untreated *T. brucei* (uninduced or induced). Dashed ellipse highlights miltefosine-mediated complementation of the Tb927.5.3560 RNAi growth defect. F) Chart summarising *T. brucei* population growth in the presence or absence of TbVAMP7B RNAi in a subset of miltefosine concentrations from five independent biological replicates. Individual growth (B) and EC_50_ (C, E) assays were carried out in triplicate and quadruplicate, respectively. Error bars represent standard deviation. *P*-values derived from paired Student’s *t*-test (** <0.01; *** <0.001).

To test this hypothesis, we generated stem-loop RNAi BSF *T. brucei* cell lines targeting TbVAMP7B and Tb927.5.3570. As predicted, depletion of Tb927.5.3570 had no effect on growth or sensitivity to amphotericin-B or miltefosine (Fig. S5). In contrast, knockdown of TbVAMP7B following induction in tetracycline at 2 ng or 1 μg.ml^−1^ resulted in a significant growth defect (Fig. 6B). To assess the contribution of TbVAMP7B to drug efficacy, we induced RNAi in 2 ng.ml^−1^ tetracycline for 24 hours and assessed drug sensitivity over a further 30 hours under inducing conditions. Incubation in low concentration tetracycline and a shorter EC_50_ analysis (as opposed to the standard 72-hour protocol) ensured that the growth defect due to TbVAMP7B RNAi knockdown was minimised, while still allowing us to test the protein’s contribution to drug action.

Unexpectedly, RNAi knockdown of TbVAMP7B reduced the amphotericin-B EC_50_, by 24% (Fig. 6C). However, TbVAMP7B depletion also resulted in a significant decrease in the Hill coefficient. Consequently, while the EC_50_ decreased upon TbVAMP7B depletion, the EC_90_ and EC_99_ increased 1.45-fold and 3-fold, respectively (Fig. 6D); the EC_25_ decreased by 44%, consistent with the effect on the EC_50_ and the change in the Hill coefficient. Therefore, small changes in TbVAMP7B expression can lead to significant loss of sensitivity to high concentration amphotericin-B, while enhancing sensitivity to the drug at low concentration. This relative resistance to high concentration amphotericin-B explains the enrichment of TbVAMP7B-targeting RNAi fragments following selection of the RNAi library in 1.5x EC_50_. In contrast, miltefosine at relatively low concentrations complemented the TbVAMP7B RNAi growth defect and further increased growth at lower concentrations (Fig. 6E, F).

Our findings indicate specific interactions between TbVAMP7B and both amphotericin-B and miltefosine. VAMP7 proteins are involved in endosome and lysosome membrane fusion (45) and it is notable in this respect that amphotericin-B disrupts membranes and miltefosine is a phospholipid drug. TbVAMP7B depletion does not significantly increase the EC_50_ for either drug but, nevertheless, these interactions may be important in a clinical setting where exposure will be variable in different tissues and at different times following dosing.

### Multiple hits link amphotericin-B action to phospholipid transport and metabolism

Our amphotericin-B screen yielded thirteen high-confidence hits, for which Gene-Ontology term profiling revealed links to membranes and lipids (Table S2; Fig. S6). This is consistent with disruption of membranes by amphotericin-B. Miltefosine uptake in *Leishmania* is dependent on a flippase (17, 18), which also contributes to the anti-leishmanial action of amphotericin-B (46). RNAi fragments targeting the syntenic locus in *T. brucei*, *Tb927.11.3350*, were enriched following selection in amphotericin-B and miltefosine (Fig. 5; Fig. 7A). Depletion of Tb927.11.3350, while having no effect on parasite growth in culture (Fig. 7B), led to a reproducible increase in amphotericin-B and miltefosine EC_50_ (Fig. 7C, D). RNAi knockdown also significantly enhanced short-term survival in high concentration amphotericin-B and miltefosine (Fig. S6). Therefore, as in *Leishmania*, the *T. brucei* miltefosine transporter orthologue contributes to the action of miltefosine and amphotericin-B.

**Figure 7.**
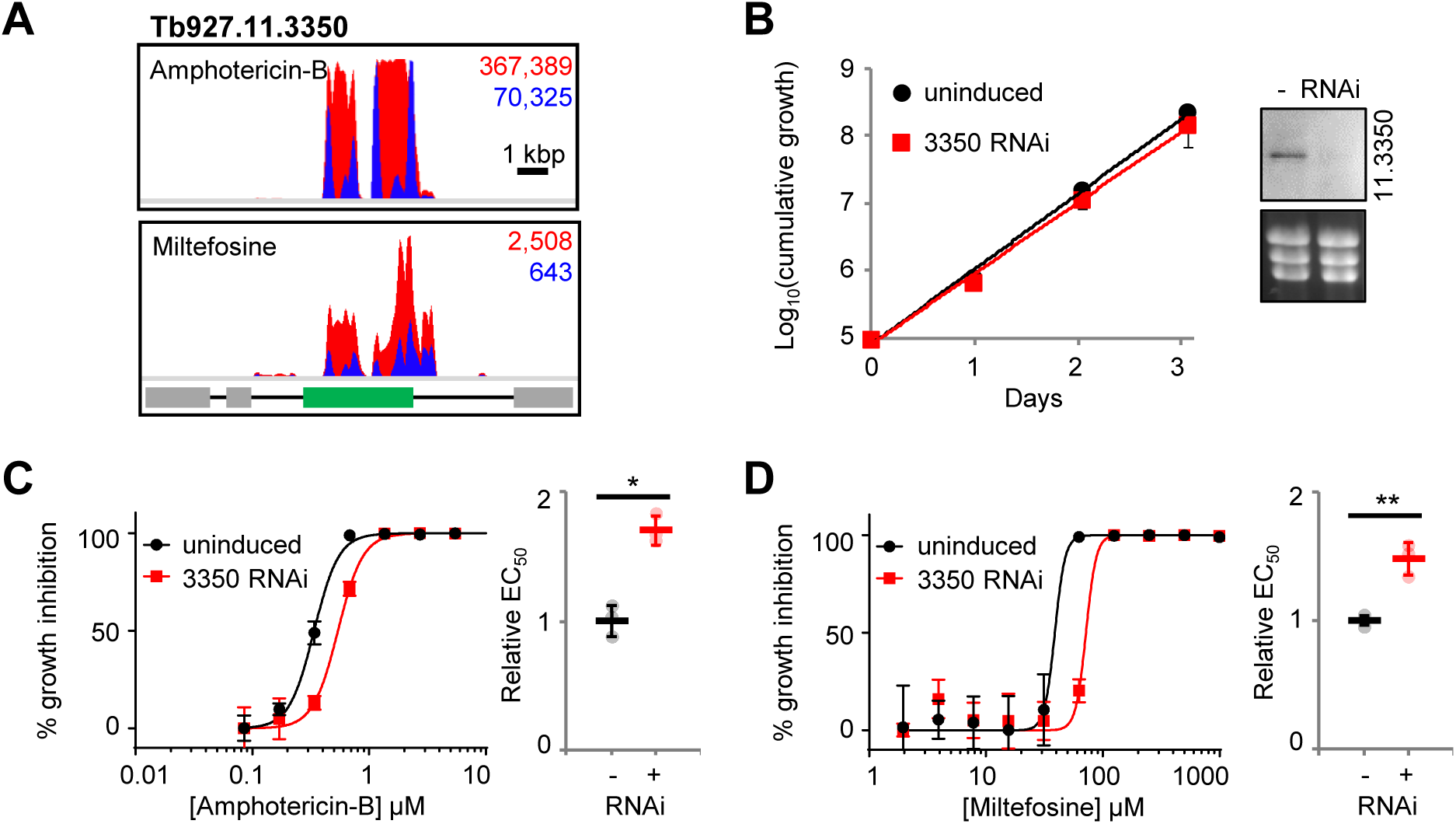
The *T. brucei* miltefosine transporter orthologue, Tb927.11.3350, influences miltefosine and amphotericin-B efficacy against *T. brucei*. A) Total (red) and RNAi construct-specific 14mer-containing (blue) reads mapping to *Tb927.11.3350* following amphotericin-B (AmB) or miltefosine selection. Targeted open reading frames highlighted in green; flanking open reading frames coloured grey. B) *T. brucei* population growth following RNAi knockdown of Tb927.11.3350. Inset: confirmation of RNAi knockdown by northern blot; ethidium bromide stained gel shown as a loading control. C, D) Representative amphotericin-B and miltefosine EC_50_ assays following RNAi knockdown of Tb927.11.3350. Inset charts summarise data from three independent biological replicates. Individual growth (B) and EC_50_ (C, D) assays were carried out in triplicate and quadruplicate, respectively. Error bars represent standard deviation. *P*-values derived from Student’s *t*-test (* <0.05; ** <0.01). RNAi inductions were carried out in 1 µg.ml^−1^ tetracycline.

In addition to Tb927.11.3350, the *T. brucei* genome contains three other putative flippases (Fig. 8A), as well as three putative β-subunits, including Tb927.11.13770, the syntenic orthologue of *Leishmania* Ros3 (18). Three of the four flippases (Tb927.4.1510, Tb927.11.3350 and Tb927.11.13000) have a similar domain organisation to the yeast flippases and possess the DEGT and DKTGT motifs characteristic of the actuator and phosphorylation domains (47). The fourth, Tb927.6.3550, lacks the flippase DEGT domain, although it clusters with the *Leishmania* flippase, LmjF.34.2630. However, it also lacks the TGES domain characteristic of the related cation transporting P-type ATPases, such as yeast Pay2 (47), so its identity is unclear (Fig. 8A).

**Figure 8.**
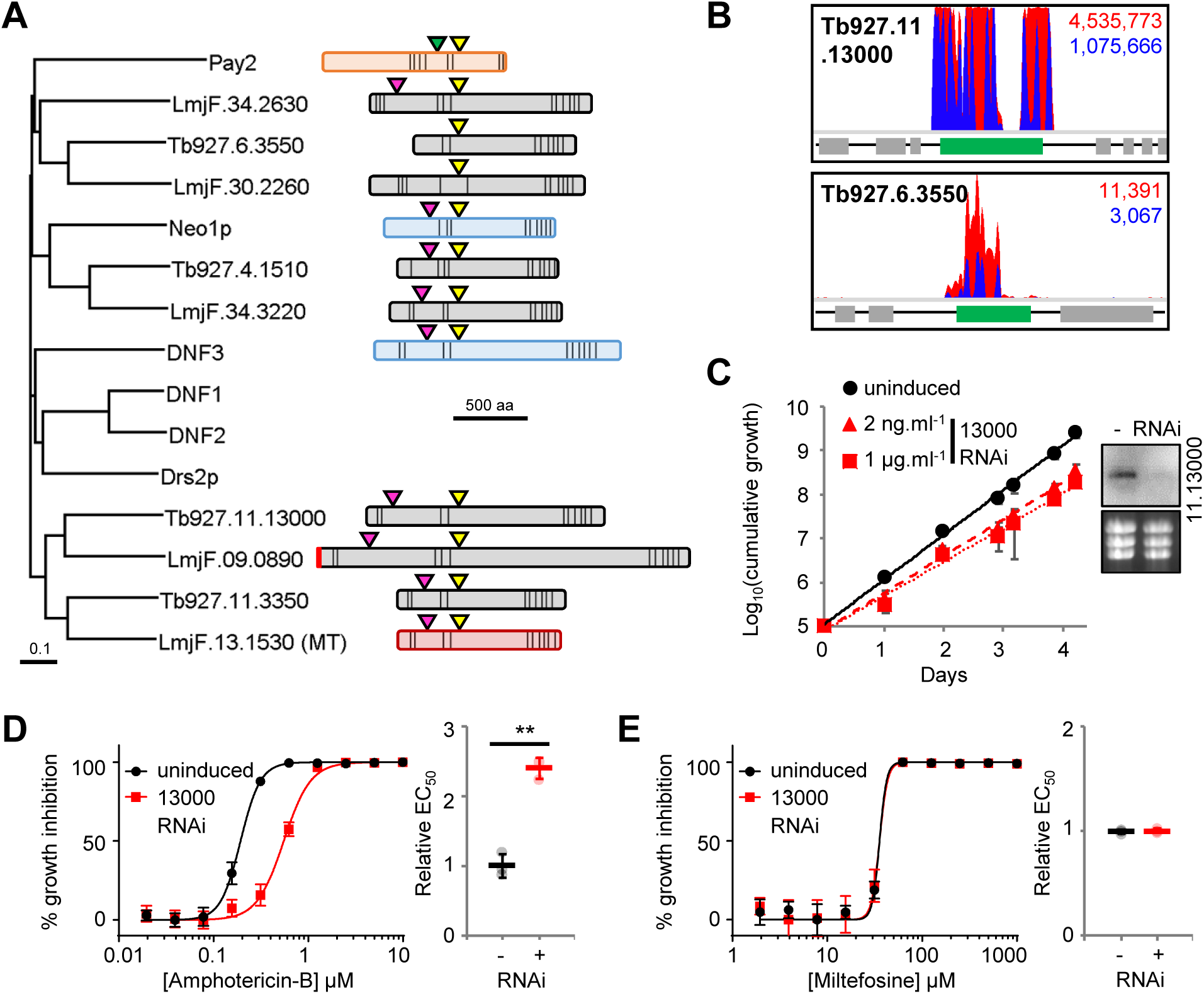
Flippases influence the action of amphotericin-B. A) Neighbour joining phylogenetic tree showing the *T. brucei* and *L. major* flippases versus the *S. cerevisiae* flippases (Neo1p, Drs2p and DNF1-3) and a representative cation-transporting P-type ATPase (Pay2). Schematics of predicted *T. brucei* and *L. major* flippases and representative *S. cerevisiae* flippases (Neo1p and DNF3) and P-type ATPase (Pay2), highlighting key conserved residues (actuator domain: TGES [green triangle], DEGT [pink triangle]; and, phosphorylation domain, DKTGT [yellow triangle]); predicted signal peptide, vertical red bar; and, predicted *trans*-membrane domain organisation, vertical black bars. B) Total (red) and RNAi construct-specific 14mer-containing (blue) reads mapping to *Tb927.11.13000* and *Tb927.6.3550* following amphotericin-B selection. Targeted open reading frames highlighted in green; flanking open reading frames coloured grey. C) *T. brucei* population growth following RNAi knockdown of Tb927.11.13000. Inset: confirmation of RNAi knockdown by northern blot; ethidium bromide stained gel shown as a loading control. D, E) Representative amphotericin-B and miltefosine EC_50_ assays following RNAi knockdown of Tb927.11.13000. Inset charts summarise data from three independent biological replicates. Individual growth (C) and EC_50_ (D, E) assays were carried out in triplicate and quadruplicate, respectively. Error bars represent standard deviation. *P*-values derived from Student’s *t*-test (* <0.05; ** <0.01). RNAi inductions were carried out in 1 µg.ml^−1^ tetracycline, unless otherwise stated.

In addition to the *Leishmania* miltefosine transporter orthologue, Tb927.11.3350, RNAi fragments targeting the flippases, Tb927.11.13000 and Tb927.6.3550, and the β-subunit, Tb927.11.13200, were enriched following selection in amphotericin-B, with Tb927.11.13000 represented by 78% of mapped reads (Fig. 8B; Table S1). Targeted RNAi depletion of Tb927.11.13000 led to a mild growth defect (Fig. 8C) and a more than two-fold EC_50_ increase, validating this protein as an amphotericin-B efficacy determinant in *T. brucei* (Fig. 8D). The impact of Tb927.11.13000 depletion was most pronounced during the initial 24 hours of drug exposure, enabling the parasite population to increase approximately 1.3-fold and four-fold over eight and 24 hours, respectively, in the presence of 0.7 µM (>EC_99_) amphotericin-B (Fig. S6). The uninduced population declined by more than 40% and 60% over the same periods. In addition, while exposure to 1.8 µM (>EC_99.9_) amphotericin-B led to an 80% decline in the induced population over 24 hours, cultures of uninduced cells were cleared within four hours exposure to this drug concentration (Fig. S6). Depletion of this putative phospholipid-transporting ATPase had no effect on miltefosine efficacy (Fig. 8E) confirming its specific contribution to amphotericin-B action.

Our results reveal that multiple *T. brucei* flippases drive the efficacy of amphotericin-B, all of which have syntenic orthologues in *Leishmania* (Fig. 8A). Therefore, in addition to the well-characterised miltefosine-transporting flippase, other *Leishmania* flippases may play significant, and potentially specific, roles in the anti-leishmanial action of amphotericin-B and miltefosine.

## Discussion

In the current absence of an effective genome-scale loss-of-function screen in *Leishmania*, we speculated that selection of a *T. brucei* RNAi library would provide insights into anti-leishmanial drug action, while also revealing novel *T. brucei* biology. By selecting our genome-scale BSF *T. brucei* RNAi library in the current anti-leishmanial drugs followed by RIT-seq analysis, we identified a panel of putative anti-leishmanial drug efficacy determinants (Table S1 and Fig. S1). SSG and miltefosine selection respectively identified TbAQP3, an orthologue of the known SSG transporter, and Tb927.11.3350, the *T. brucei* orthologue of the *Leishmania* miltefosine transporter, confirming the power of this approach. In addition to these known drug transporters, we validated several novel drug efficacy determinants identified by our selective screens: Tb927.9.6360-80 (paromomycin), Tb927.5.3560 (miltefosine and amphotericin-B) and Tb927.11.13000 (amphotericin-B). Our results highlight the role of a lysosomal transporter in paromomycin efficacy, emphasise the importance of membrane composition in the action of amphotericin-B and miltefosine, provide insight into the substrate selectivity of the trypanosomatid aquaglyceroporins, and present several new candidate anti-leishmanial drug efficacy determinants (Fig. 9).

**Figure 9.**
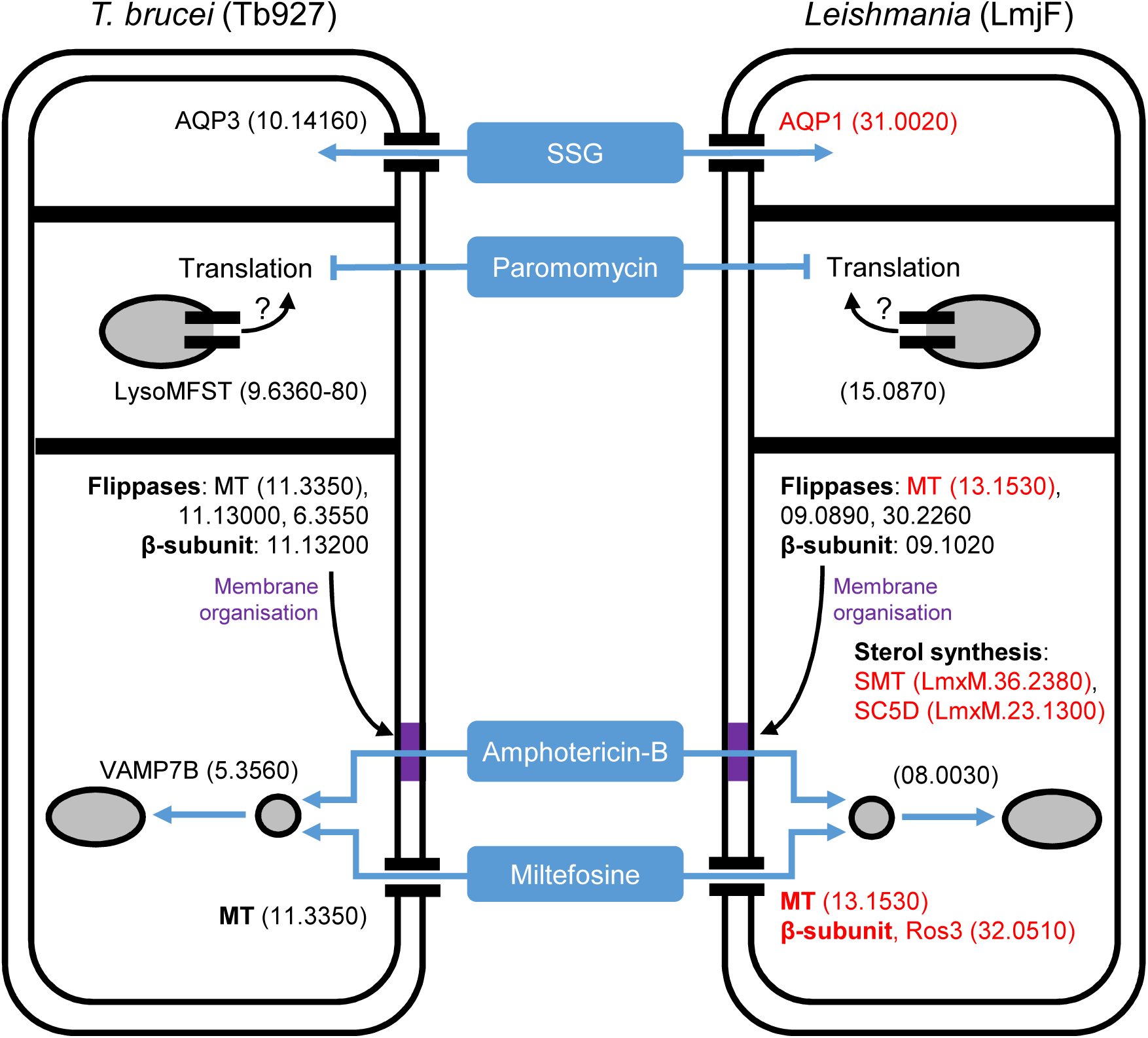
Known and candidate drivers of anti-leishmanial drug efficacy in *Leishmania*. The key *T. brucei* proteins identified in our anti-leishmanial loss-of-function screen (left hand panel) and their *Leishmania* orthologues (right hand panel) represent candidate anti-leishmanial drug efficacy determinants. Red denotes known *Leishmania* drivers of anti-leishmanial efficacy whose loss-of-function reduces drug efficacy (see text for details). The strain prefix for the truncated gene IDs is at the top of each panel, with the exception of the sterol biosynthetic enzymes recently shown to contribute to amphotericin-B efficacy against *L. mexicana* (70). Grey-filled circles (endosomes) and ellipses (lysosome) represent the endocytic system. The purple block represents membrane modified by changes in sterol biosynthesis and the putative action of the flippases and their β-subunit; changes in membrane composition anywhere in the endocytic system may influence the intracellular transit of amphotericin-B or its ability to form ion permeable channels.

*T. brucei* RNAi library selection in SSG and our subsequent validation experiments identified a single efficacy determinant, TbAQP3. Aquaglyceroporins are ubiquitous transporters of water, glycerol and other small solutes, whose specificity is defined by their selectivity filter residues. *Leishmania* AQP1 and the *T. brucei* proteins, TbAQP1 and TbAQP3, have the same selectivity filter, NPA/NPA/WGYR, while TbAQP2 possesses a divergent filter, NSA/NPS/IVLL (39). TbAQP2 is a key drug transporter in *T. brucei*, mediating the uptake of pentamidine and melarsoprol, and its loss contributes to clinical drug resistance (35-37). In addition, TbAQP2 plays an important role in glycerol transport, as its loss increases parasite sensitivity to alternative oxidase inhibition, which leads to elevated intracellular glycerol levels (48). The *in vivo* roles of the other *T. brucei* aquaglyceroporins remain unknown, though all three are capable of arsenite and antimonite transport in yeast and *Xenopus* heterologous expression systems (49). In contrast, our data demonstrate that in *T. brucei* these transporters are selective for arsenic-containing melarsoprol (TbAQP2; (35)) and antimony-containing, SSG (TbAQP3). Intriguingly, RNAi library selection with SSG failed to identify TbAQP1 even though it contains the same selectivity filter as TbAQP3. This suggests important functional and regulatory differences between TbAQP1 and TbAQP3, which may influence their ability to contribute to SSG uptake in bloodstream-form *T. brucei*. For example, TbAQP3 is localised to the plasma membrane in bloodstream-form *T. brucei* and TbAQP1 localises to the flagella membrane (35, 50). This differential localisation may influence their ability to mediate antimonial uptake.

The aminoglycoside, paromomycin, is thought to inhibit protein synthesis in *Leishmania* and enters the cell *via* endocytosis (21, 51, 52). However, RNAi library selection did not identify a surface receptor suggesting that, at least in *T. brucei*, paromomycin entry is not dependent on a specific ligand-receptor interaction. Rather, the high endocytic flux associated with surface VSG internalisation (53) may drive drug uptake. RNAi fragments targeting Tb927.9.6360-80 dominated the paromomycin-selected RNAi library, with the remaining 28 high confidence hits constituting only 9% of mapped reads. This locus encodes a set of closely related MFST proteins, at least one of which localises to the lysosome, and has previously been associated with suramin efficacy (24). In contrast to paromomycin, several other endocytic pathway proteins, including three lysosomal proteins (p67, cathepsin-L and the MFST proteins), influence suramin efficacy (24). This led to the proposal that proteolytic processing in the lysosome releases suramin from bound proteins, enabling neutralisation in the acidic environment or association with an alternative endogenous carrier and escape to the cytoplasm *via* one or more of the lysosomal MFSTs (54). In contrast, the absence of hits targeting other endocytic components following paromomycin RNAi library selection suggests little reliance on the endocytic network *per se*. Therefore, the lysosomal MFST proteins may influence paromomycin efficacy indirectly. MFST proteins mediate the transit of a diverse range of molecules, including polyamines and amino acids (41), and changes in the intracellular flux of these molecules may affect translation efficiency, which in turn may influence paromomycin efficacy. Deletion of the *Tb927.9.6360-80* locus from *T. brucei* yields only a two-fold increase in paromomycin EC_50_. However, the MFST protein encoded by the syntenic single copy gene in *Leishmania* (e.g. *LmjF.15.0870*) remains to be characterised and may make a more substantial contribution to paromomycin action against this parasite.

Combination therapies are increasingly being used to treat leishmaniasis, enabling reduced dosing and treatment duration, resulting in fewer side effects (8). For example, a single dose of liposomal amphotericin-B in combination with a short course of oral miltefosine or intramuscular paromomycin is an effective treatment for visceral leishmaniasis (VL) in the Indian sub-continent (55). In East Africa, SSG-paromomycin combination therapy is effective against VL (56). However, *L. donovani* resistant to these and other anti-leishmanial drug combinations can be selected for *in vitro* (9, 10), and oxidative defence upregulation and changes in membrane fluidity have been associated with cross-resistance in laboratory-derived lines (23). Therefore, we carried out pairwise comparisons of our RNAi library screen data to identify potential cross-efficacy determinants. Only two hits fulfilled our stringency criteria, both of which influence amphotericin-B and miltefosine action: TbVAMP7B, an endosomal SNARE protein responsible for endosomal-lysosomal fusion in other eukaryotes (45, 57), and Tb927.11.3350, the *T. brucei* orthologue of the *Leishmania* miltefosine transporter (17). However, while both of these proteins may influence membrane fluidity (see below), it seems unlikely that either contributes significantly to oxidative defence. Recent Cos-seq gain-of-function analyses in *L. infantum* identified several candidate proteins whose overexpression reduces sensitivity to multi-drug exposure (26); these also lack an obvious connection to oxidative defence. Therefore, rather than being dependent on the increase or decrease in expression of a single protein, changes in oxidative defence that lead to anti-leishmanial resistance are likely to be multi-factorial. Our findings also suggest that miltefosine/amphotericin-B combination therapy is the most vulnerable to loss-of-function mutation, while others may be less susceptible to the down-regulation of a single protein. This finding is particularly significant, given that recent trials have confirmed the efficacy of amphotericin-B/miltefosine combination therapy in treating VL (58, 59).

In contrast to the other anti-leishmanial drug efficacy determinants described herein, TbVAMP7B depletion does not simply increase the drugs’ EC_50_. Instead, TbVAMP7B RNAi knockdown reduces amphotericin-B EC_50_ and has little effect on miltefosine EC_50_. The drop in amphotericin-B EC_50_ is due to a substantial decrease in the amphotericin-B Hill coefficient, which has the opposite effect on EC_90_ and EC_99_, increasing both and enabling TbVAMP7B-depleted parasites to persist at these drug concentrations. Our data shows that *T. brucei* has limited tolerance for TbVAMP7B depletion, presumably due to impairment of endosomal-lysosomal fusion (45). Intriguingly, exposure to low concentration miltefosine complements the growth defect seen following TbVAMP7B depletion, suggesting that miltefosine treatment is able to promote vesicle membrane fusion in the endocytic system, a possible consequence of the enhanced membrane fluidity seen upon miltefosine exposure (60). TbVAMP7B has also recently been identified as a putative *T. brucei* apolipoprotein-L1 sensitivity determinant (61), and other workers have highlighted the importance of the intracellular transit of apoL1-carrying membrane to trypanolysis (62, 63). Our findings suggest that such transit also contributes to amphotericin-B and miltefosine action. The VAMP7 proteins are highly conserved between *T. brucei* and *Leishmania* (43), suggesting that *Leishmania* parasites will also be sensitive to VAMP7B loss (LmjF.08.0030). However, subtle changes in VAMP7B expression that can be tolerated may enable parasites to take advantage of variations in amphotericin-B and miltefosine tissue penetrance.

Miltefosine uptake in *Leishmania* is dependent on a phospholipid-transporting flippase (the miltefosine transporter, MT) and its β-subunit, Ros3 (17, 18); both *in vitro* selected lines and miltefosine resistant *L. donovani* clinical isolates harbour mutations in the MT (7, 64, 65). Consistent with this, *T. brucei* RNAi library selection in miltefosine led to enrichment for RNAi fragments mapping to the syntenic sequence in *T. brucei* (*Tb927.11.3350*). RNAi library selection in amphotericin-B also enriched for RNAi fragments mapping to this gene, consistent with recent findings in *Leishmania* (46), as well as two other flippases and a putative β-subunit (Tb927.11.13200). Interestingly, the β-subunit targeted was not the syntenic orthologue of Ros3, previously shown to interact with the MT (18). Therefore, different flippase/β-subunit dependencies may have evolved following divergence of the *Leishmania* and *T. brucei* lineages. A further difference in the behaviour of these proteins between *Leishmania* and *T. brucei* lies in their localisation. The MT and Ros3 localise to the plasma membrane in *Leishmania* (18), whereas the *T. brucei* MT orthologue (Tb927.11.3350) and a second flippase (Tb927.11.13000) localise to an intracellular structure reminiscent of the endosomal system in procyclic form *T. brucei* (www.TrypTag.org; (66)). Therefore, while flippases influence drug action against *Leishmania* and *T. brucei*, they may mediate drug and/or phospholipid transit across different membranes in each parasite.

Phospholipid transport by flippases maintains the membrane asymmetry necessary for membrane fusion, vesicle trafficking and sterol homeostasis (47). The identification of a single flippase following miltefosine selection is consistent with its role as a drug transporter (17). In contrast, amphotericin-B selection identified three flippases, suggesting an indirect role in drug action, possibly through changes in membrane composition and transit through the endosomal system (Fig. 9). Amphotericin-B acts by binding membrane ergosterol (67), leading to the formation of ion-permeable channels and downstream oxidative damage (68). Consistent with the importance of ergosterol to amphotericin-B action, resistant clinical isolates exhibit elevated membrane fluidity and reduced ergosterol content (69). Recent findings have highlighted the loss of key sterol biosynthetic enzymes, and reduced ergosterol production, as a driver of resistance in laboratory-derived amphotericin-B resistant *L. mexicana* (70). Changes in flippase expression may similarly affect ergosterol membrane content and/or accessibility, thereby reducing sensitivity to amphotericin-B. Therefore, functional characterisation of the syntenic orthologues of these proteins in *Leishmania* may provide further insights into the processes and factors that drive the anti-leishmanial action of amphotericin-B.

In summary, using our genome-scale BSF *T. brucei* RNAi library we have identified a panel of putative anti-leishmanial drug efficacy determinants, highlighting two candidate cross-efficacy determinants, as well as roles for multiple flippases in the action of amphotericin-B. The findings from this orthology-based chemogenomic profiling approach substantially advance our understanding of anti-leishmanial drug mode-of-action and potential resistance mechanisms, and should facilitate the development of improved therapies, as well as surveillance for drug-resistant parasites.

## Methods

### *T. brucei* strains

MITat1.2/2T1 BSF *T. brucei* (71) were maintained in HMI11 (Invitrogen, LifeTech) supplemented with 10% foetal calf serum (Sigma) at 37°C/5% CO_2_. Transfection was carried out in either cytomix or Tb-BSF buffer (72), for integration at the 2T1 ‘landing pad’ (71, 73) or *Tb927.9.6360-80*, respectively, using a Nucleofector (Lonza) set to programme X-001. Transformants were selected in 2.5 µg.ml^−1^ hygromycin, 2 µg.ml^−1^ puromycin or 10 µg.ml^−1^ blasticidin, as appropriate. The BSF *T. brucei* RNAi library was maintained in 1 µg.ml^−1^ phleomycin and 5 µg.ml^−1^ blasticidin (34). For growth assays, cultured BSF *T. brucei* were seeded at ∼10^5^ cells.ml^−1^, counted using a haemocytometer, and diluted back every 24 hours, as necessary, for three days in the absence of antibiotics. All selective antibiotics were purchased from Invivogen.

### Drug sensitivity assays

Half-maximal effective concentrations (EC_50_) of the anti-leishmanial drugs (sodium stibogluconate, GSK; paromomycin, Sigma; miltefosine, Paladin; amphotericin-B, E R Squibb, UK) and neomycin (G418, Invivogen) were determined over 78 or 30 hours. BSF *T. brucei* were seeded at 2×10^3^ (or 2×10^5^) cells.ml^−1^ in 96-well plates in a 2-fold dilution series of each drug; assays were carried out in the absence of other antibiotics. After 72 or 24 hours, resazurin (Sigma) in PBS was added to a final concentration of 12.5 µg.ml^−1^ per well, and the plates incubated for a further 6 hours at 37°C. Fluorescence was determined using a fluorescence plate reader (Molecular Devices) at an excitation wavelength of 530 nm, an emission wavelength of 585 nm and a filter cut-off of 570 nm (74). Data were processed in Microsoft Excel, and non-linear regression analysis carried out in GraphPad Prism. The short-term kinetics of killing in high concentration drug (>EC_99_) were determined in triplicate over 24 hours from a starting cell density of 1×10^5^ cells.ml^−1^.

### *T. brucei* RNAi library screening and RIT-seq

RNA library screening was carried out as previously described (34). Briefly, library expression was induced in 1 µg.ml^−1^ tetracycline (Sigma) for 24 hours prior to selection in each anti-leishmanial drug at 1-3X EC_50_. Cell density was assessed daily using a haemocytometer and diluted to no less than 20 million cells in 100 ml media; induction and anti-leishmanial drug selection were maintained throughout. Once robust growth had been achieved for at least two days, genomic DNA was prepared for RNAi target identification. The RNAi cassettes remaining in the anti-leishmanial-selected RNAi libraries were amplified from genomic DNA using the LIB2F/LIB2R primers and sequenced on an Illumina HiSeq platform at the Beijing Genome Institute.

The sequenced RNAi target fragments were mapped against the *T. brucei strain* TREU927 reference genome (release 6.0), as described (34). Briefly, mapping was carried out using Bowtie2 (75) set to ‘very sensitive local’ alignment and output SAM files were processed using SAMtools (76). The resultant BAM files were viewed against the reference genome in the Artemis genome browser (77). Reads containing the RNAi construct-specific 14-base barcode were identified using a custom script (34), and corresponded to at least 22% of reads from each selected RNAi library. This subset of reads were mapped against the TREU927 reference genome, as above. Plots were generated using the Artemis graph tool and processed in Adobe Photoshop Elements 8.0. Stacks of reads that included the 14-base barcode on the positive strand were used to define RNAi target fragment junctions and to assign high-confidence hits as those identified by at least two RNAi target fragments. RNAi target fragment read numbers were converted to RPKM (reads/kilobase/million reads mapped) to account for inter-library read-depth variations when comparing RNAi library sequencing outputs.

Alignments were carried out in Clustal Omega (https://www.ebi.ac.uk/Tools/msa/clustalo/), unrooted neighbour joining trees were formatted in Dendroscope 3 (http://dendroscope.org/) (78) and putative *trans*-membrane domains identified using TOPCONS (http://topcons.cbr.su.se/) (79). GO-term profiles were constructed using the GO analysis tool at http://tritrypdb.org.

### Plasmid and *T. brucei* strain construction and analysis

*Tb927.9.6360-80* locus targeting fragments were cloned into pPAC and pBSD, enabling replacement of both alleles of the three-gene locus with puromycin acetyltransferase (*PAC*) and blasticidin-S deaminase (*BSD*) open reading frames. Stem-loop RNAi constructs targeting Tb927. 11.6680 (AAT15), Tb927.11.13000, Tb927.11.3350, Tb927.5.3560 (TbVAMP7B) and Tb927.5.3570 were assembled in pRPa-iSL (73). RNAi targeting fragments were designed using the RNAit primer design algorithm to minimise off-target effects (80). pRPa-iSL constructs were linearised with *Asc*I (NEB) prior to transfection and targeted integration at the *rDNA* spacer ‘landing pad’ locus in 2T1 BSF *T. brucei* (71). Details of all primers are available upon request. Tb927.9.6360-80 allelic replacement was confirmed by Southern hybridisation following *Xho*I (New England Biolabs) digestion of genomic DNA. RNAi knockdown was confirmed by northern hybridisation of total RNA or, in the case of Tb927.11.6680, by RT-qPCR, as described (81). For Southern and northern hybridisation, digoxigenin-dUTP (Roche) labelled DNA probes were generated by PCR, hybridised and detected according to standard protocols and the manufacturer’s instructions.

### Data availability

Sequence data are available as fastq files at the European Nucleotide Archive (https://www.ebi.ac.uk/ena) under study accession number, PRJEB31973 (amphotericin-B, ERS3348616; miltefosine, ERS3348617; paromomycin, ERS3348618; sodium stibogluconate, ERS3348619).

## Figure legends

**Figure S1**

**Candidate anti-leishmanial drug efficacy determinants identified by *T. brucei* RNAi library selection**. Total (red) and RNAi construct-specific 14mer-containing (blue) reads mapping to individual loci following BSF *T. brucei* RNA library selection in paromomycin (A), amphotericin-B (B) and miltefosine (C). Targeted open reading frames highlighted in green; flanking open reading frames coloured grey. Where a substantial number of reads target regions outside the open reading frame, the predicted untranslated region is highlighted by a narrow green bar. See Table S1 for further details.

**Figure S2**

**Neither AAT15 (Tb927.11.6680) depletion nor Tudor Staphylococcal nuclease (Tb927.11.14190) deletion affects aminoglycoside efficacy against BSF *T. brucei* over 72 hours.** A, B) Total (red) and RNAi construct-specific 14mer-containing (blue) reads mapping to *Tb927.11.*6680 (A) and *Tb927.11.14190* (B) following paromomycin selection. Targeted open reading frames highlighted in green; flanking open reading frames coloured grey. C) *T. brucei* population growth following AAT15 RNAi knockdown. Inset: RNA depletion was confirmed by RT-qPCR following 24-hour induction in 1 µg.ml^−1^ tetracycline. D, E) Representative paromomycin and neomycin EC_50_ assays following AAT15 RNAi knockdown induced in 1 µg.ml^−1^ tetracycline. F, G) Representative paromomycin (D) and neomycin (E) EC_50_ assays comparing wild type and *Tb927.11.14190* null (*tsn*) BSF *T. brucei*. Inset charts summarise data from three independent biological replicates. Individual growth (C) and EC_50_ (D-G) assays were carried out in triplicate and quadruplicate, respectively. Error bars represent standard deviation.

**Figure S3**

***Tb927.9.6360-80* clusters with the syntenic *LmjF.15.0870.*** Twenty nine open reading frames annotated ‘major facilitator’ or ‘MFS’ in the L. major Friedlin reference genome were aligned with the Tb927.9.6360-80 open reading frames using Clustal Omega (https://www.ebi.ac.uk/Tools/msa/clustalo/). The unrooted neighbour joining phylogenetic tree was formatted in Dendroscope 3 (http://dendroscope.org/).

**Figure S4**

***MFST* locus null *T. brucei* exhibit enhanced tolerance to high concentration aminoglycosides.** Relative population growth of wild type (WT) and *MFST* locus null (*mfst*) *T. brucei* in A) paromomycin and B) neomycin at >EC_99_. Assays were carried out in triplicate. Error bars represent standard deviation.

**Figure S5**

**Tb927.5.3570 does not contribute to the efficacy of amphotericin-B or miltefosine against *T. brucei*.** A) *T. brucei* population growth following Tb927.5.3570 RNAi knockdown. Inset: confirmation of RNAi knockdown by northern blot; ethidium bromide stained gel shown as a loading control. B, C) Representative amphotericin-B and miltefosine EC_50_ assays following Tb927.5.3570 RNAi knockdown. RNAi inductions were carried out in 1 µg.ml^−1^ tetracycline. Individual growth (A) and EC_50_ (B, C) assays were carried out in triplicate and quadruplicate, respectively. Error bars represent standard deviation.

**Figure S6**

**Gene Ontology analysis of the high confidence hits identified following amphotericin-B RNAi library selection.** Plot generated using the GO analysis tool at TritrypDB.org. Point diameter corresponds to relative number of proteins in each category. See Table S2 for further details.

**Figure S7**

***T. brucei* exhibit enhanced tolerance to high concentration amphotericin-B following flippase depletion.** A, C) Representative assays showing relative population growth in >EC_99_ amphotericin-B following (A) Tb927.11.3350 and (C) Tb927.11.13000 RNAi knockdown. B, D) Relative population growth in >EC_99_ amphotericin-B following (B) Tb927.11.3350 and (D) Tb927.11.13000 RNAi knockdown; data derived from three independent biological replicates. Individual growth assays were carried out in triplicate. Error bars represent standard deviation. *P*-values derived from Student’s *t*-test (* <0.05; *** <0.001). RNAi inductions were carried out in 1 µg.ml^−1^ tetracycline.

**Table S1**

Transcripts represented by >99 RNAi construct-specific barcode-containing reads per kilobase per transcript following BSF *T. brucei* RNAi library selection in the anti-leishmanial drugs.

**Table S2**

Gene Ontology analysis of the high confidence hits identified by amphotericin-B RNAi library selection.

## Acknowledgements

This work was funded by a Wellcome Trust Institutional Strategic Support Fund (LSHTM) fellowship (www.wellcome.ac.uk/) awarded to SA. DH is a Wellcome Trust Investigator award recipient (100320/Z/12/Z). HBSS was supported by the Biotechnology and Biological Sciences Research Council (BB/J014567/1). MVS was supported by the Leonardo da Vinci internship programme. The funders had no role in study design, data collection and interpretation, or the decision to submit the work for publication. Thanks to Dr Vanessa Yardley, LSHTM, for sharing her stocks of the anti-leishmanial drugs, sodium stibogluconate, miltefosine and amphotericin-B. Thanks to the ‘Advanced training in molecular biology’ (LSHTM) class of 2017 for the *MFST* null Southern images.

## References

1. Torres-Guerrero E, Quintanilla-Cedillo MR, Ruiz-Esmenjaud J, Arenas R. 2017. Leishmaniasis: a review. F1000Res 6:750.

2. Buscher P, Cecchi G, Jamonneau V, Priotto G. 2017. Human African trypanosomiasis. Lancet 390:2397–2409.

3. Perez-Molina JA, Molina I. 2018. Chagas disease. Lancet 391:82–94.

4. Steverding D. 2017. The history of leishmaniasis. Parasit Vectors 10:82.

5. Barrett MP, Croft SL. 2012. Management of trypanosomiasis and leishmaniasis. Br Med Bull 104:175–96.

6. Mandal S, Maharjan M, Singh S, Chatterjee M, Madhubala R. 2010. Assessing aquaglyceroporin gene status and expression profile in antimony-susceptible and -resistant clinical isolates of *Leishmania donovani* from India. J Antimicrob Chemother 65:496–507.

7. Srivastava S, Mishra J, Gupta AK, Singh A, Shankar P, Singh S. 2017. Laboratory confirmed miltefosine resistant cases of visceral leishmaniasis from India. Parasit Vectors 10:49.

8. Ponte-Sucre A, Gamarro F, Dujardin JC, Barrett MP, Lopez-Velez R, Garcia-Hernandez R, Pountain AW, Mwenechanya R, Papadopoulou B. 2017. Drug resistance and treatment failure in leishmaniasis: A 21st century challenge. PLoS Negl Trop Dis 11:e0006052.

9. Garcia-Hernandez R, Manzano JI, Castanys S, Gamarro F. 2012. *Leishmania donovani* develops resistance to drug combinations. PLoS Negl Trop Dis 6:e1974.

10. Hendrickx S, Inocencio da Luz RA, Bhandari V, Kuypers K, Shaw CD, Lonchamp J, Salotra P, Carter K, Sundar S, Rijal S, Dujardin JC, Cos P, Maes L. 2012. Experimental induction of paromomycin resistance in antimony-resistant strains of *L. donovani*: outcome dependent on in vitro selection protocol. PLoS Negl Trop Dis 6:e1664.

11. Mesu V, Kalonji WM, Bardonneau C, Mordt OV, Blesson S, Simon F, Delhomme S, Bernhard S, Kuziena W, Lubaki JF, Vuvu SL, Ngima PN, Mbembo HM, Ilunga M, Bonama AK, Heradi JA, Solomo JLL, Mandula G, Badibabi LK, Dama FR, Lukula PK, Tete DN, Lumbala C, Scherrer B, Strub-Wourgaft N, Tarral A. 2018. Oral fexinidazole for late-stage African *Trypanosoma brucei gambiense* trypanosomiasis: a pivotal multicentre, randomised, non-inferiority trial. Lancet 391:144–154.

12. Wyllie S, Patterson S, Stojanovski L, Simeons FR, Norval S, Kime R, Read KD, Fairlamb AH. 2012. The anti-trypanosome drug fexinidazole shows potential for treating visceral leishmaniasis. Sci Transl Med 4:119re1.

13. Deeks ED. 2019. Fexinidazole: First Global Approval. Drugs 79:215–220.

14. Hendrickx S, Guerin PJ, Caljon G, Croft SL, Maes L. 2018. Evaluating drug resistance in visceral leishmaniasis: the challenges. Parasitology 145:453–463.

15. Downing T, Imamura H, Decuypere S, Clark TG, Coombs GH, Cotton JA, Hilley JD, de Doncker S, Maes I, Mottram JC, Quail MA, Rijal S, Sanders M, Schonian G, Stark O, Sundar S, Vanaerschot M, Hertz-Fowler C, Dujardin JC, Berriman M. 2011. Whole genome sequencing of multiple Leishmania donovani clinical isolates provides insights into population structure and mechanisms of drug resistance. Genome Res 21:2143–56.

16. Rastrojo A, Garcia-Hernandez R, Vargas P, Camacho E, Corvo L, Imamura H, Dujardin JC, Castanys S, Aguado B, Gamarro F, Requena JM. 2018. Genomic and transcriptomic alterations in Leishmania donovani lines experimentally resistant to antileishmanial drugs. Int J Parasitol Drugs Drug Resist 8:246–264.

17. Perez-Victoria FJ, Gamarro F, Ouellette M, Castanys S. 2003. Functional cloning of the miltefosine transporter. A novel P-type phospholipid translocase from Leishmania involved in drug resistance. J Biol Chem 278:49965–71.

18. Perez-Victoria FJ, Sanchez-Canete MP, Castanys S, Gamarro F. 2006. Phospholipid translocation and miltefosine potency require both *L. donovani* miltefosine transporter and the new protein LdRos3 in *Leishmania* parasites. J Biol Chem 281:23766–75.

19. Gourbal B, Sonuc N, Bhattacharjee H, Legare D, Sundar S, Ouellette M, Rosen BP, Mukhopadhyay R. 2004. Drug uptake and modulation of drug resistance in *Leishmania* by an aquaglyceroporin. J Biol Chem 279:31010–7.

20. El Fadili K, Messier N, Leprohon P, Roy G, Guimond C, Trudel N, Saravia NG, Papadopoulou B, Legare D, Ouellette M. 2005. Role of the ABC transporter MRPA (PGPA) in antimony resistance in Leishmania infantum axenic and intracellular amastigotes. Antimicrob Agents Chemother 49:1988–93.

21. Chawla B, Jhingran A, Panigrahi A, Stuart KD, Madhubala R. 2011. Paromomycin affects translation and vesicle-mediated trafficking as revealed by proteomics of paromomycin - susceptible -resistant *Leishmania donovani*. PLoS One 6:e26660.

22. Brotherton MC, Bourassa S, Legare D, Poirier GG, Droit A, Ouellette M. 2014. Quantitative proteomic analysis of amphotericin B resistance in *Leishmania infantum*. Int J Parasitol Drugs Drug Resist 4:126–32.

23. Berg M, Garcia-Hernandez R, Cuypers B, Vanaerschot M, Manzano JI, Poveda JA, Ferragut JA, Castanys S, Dujardin JC, Gamarro F. 2015. Experimental resistance to drug combinations in Leishmania donovani: metabolic and phenotypic adaptations. Antimicrob Agents Chemother 59:2242–55.

24. Alsford S, Eckert S, Baker N, Glover L, Sanchez-Flores A, Leung KF, Turner DJ, Field MC, Berriman M, Horn D. 2012. High-throughput decoding of antitrypanosomal drug efficacy and resistance. Nature 482:232–6.

25. Alsford S, Kelly JM, Baker N, Horn D. 2013. Genetic dissection of drug resistance in trypanosomes. Parasitology 140:1478–91.

26. Gazanion E, Fernandez-Prada C, Papadopoulou B, Leprohon P, Ouellette M. 2016. Cos-Seq for high-throughput identification of drug target and resistance mechanisms in the protozoan parasite *Leishmania*. Proc Natl Acad Sci U S A 113:E3012–21.

27. Corpas-Lopez V, Moniz S, Thomas M, Wall RJ, Torrie LS, Zander-Dinse D, Tinti M, Brand S, Stojanovski L, Manthri S, Hallyburton I, Zuccotto F, Wyatt PG, De Rycker M, Horn D, Ferguson MAJ, Clos J, Read KD, Fairlamb AH, Gilbert IH, Wyllie S. 2018. Pharmacological Validation of N-Myristoyltransferase as a Drug Target in Leishmania donovani. ACS Infect Dis doi:10.1021/acsinfecdis.8b00226.

28. Fernandez-Prada C, Sharma M, Plourde M, Bresson E, Roy G, Leprohon P, Ouellette M. 2018. High-throughput Cos-Seq screen with intracellular *Leishmania infantum* for the discovery of novel drug-resistance mechanisms. Int J Parasitol Drugs Drug Resist 8:165–173.

29. Lye LF, Owens K, Shi H, Murta SM, Vieira AC, Turco SJ, Tschudi C, Ullu E, Beverley SM. 2010. Retention and loss of RNA interference pathways in trypanosomatid protozoans. PLoS Pathog 6:e1001161.

30. El-Sayed NM, Myler PJ, Blandin G, Berriman M, Crabtree J, Aggarwal G, Caler E, Renauld H, Worthey EA, Hertz-Fowler C, Ghedin E, Peacock C, Bartholomeu DC, Haas BJ, Tran AN, Wortman JR, Alsmark UC, Angiuoli S, Anupama A, Badger J, Bringaud F, Cadag E, Carlton JM, Cerqueira GC, Creasy T, Delcher AL, Djikeng A, Embley TM, Hauser C, Ivens AC, Kummerfeld SK, Pereira-Leal JB, Nilsson D, Peterson J, Salzberg SL, Shallom J, Silva JC, Sundaram J, Westenberger S, White O, Melville SE, Donelson JE, Andersson B, Stuart KD, Hall N. 2005. Comparative genomics of trypanosomatid parasitic protozoa. Science 309:404–9.

31. Khare S, Nagle AS, Biggart A, Lai YH, Liang F, Davis LC, Barnes SW, Mathison CJ, Myburgh E, Gao MY, Gillespie JR, Liu X, Tan JL, Stinson M, Rivera IC, Ballard J, Yeh V, Groessl T, Federe G, Koh HX, Venable JD, Bursulaya B, Shapiro M, Mishra PK, Spraggon G, Brock A, Mottram JC, Buckner FS, Rao SP, Wen BG, Walker JR, Tuntland T, Molteni V, Glynne RJ, Supek F. 2016. Proteasome inhibition for treatment of leishmaniasis, Chagas disease and sleeping sickness. Nature 537:229–233.

32. Seifert K, Escobar P, Croft SL. 2010. In vitro activity of anti-leishmanial drugs against Leishmania donovani is host cell dependent. J Antimicrob Chemother 65:508–11.

33. Berriman M, Ghedin E, Hertz-Fowler C, Blandin G, Renauld H, Bartholomeu DC, Lennard NJ, Caler E, Hamlin NE, Haas B, Bohme U, Hannick L, Aslett MA, Shallom J, Marcello L, Hou L, Wickstead B, Alsmark UC, Arrowsmith C, Atkin RJ, Barron AJ, Bringaud F, Brooks K, Carrington M, Cherevach I, Chillingworth TJ, Churcher C, Clark LN, Corton CH, Cronin A, Davies RM, Doggett J, Djikeng A, Feldblyum T, Field MC, Fraser A, Goodhead I, Hance Z, Harper D, Harris BR, Hauser H, Hostetler J, Ivens A, Jagels K, Johnson D, Johnson J, Jones K, Kerhornou AX, Koo H, Larke N, et al. 2005. The genome of the African trypanosome *Trypanosoma brucei*. Science 309:416–22.

34. Glover L, Alsford S, Baker N, Turner DJ, Sanchez-Flores A, Hutchinson S, Hertz-Fowler C, Berriman M, Horn D. 2015. Genome-scale RNAi screens for high-throughput phenotyping in bloodstream-form African trypanosomes. Nat Protoc 10:106–33.

35. Baker N, Glover L, Munday JC, Aguinaga Andres D, Barrett MP, de Koning HP, Horn D. 2012. Aquaglyceroporin 2 controls susceptibility to melarsoprol and pentamidine in African trypanosomes. Proc Natl Acad Sci U S A 109:10996–1001.

36. Song J, Baker N, Rothert M, Henke B, Jeacock L, Horn D, Beitz E. 2016. Pentamidine Is Not a Permeant but a Nanomolar Inhibitor of the *Trypanosoma brucei* Aquaglyceroporin-2. PLoS Pathog 12:e1005436.

37. Graf FE, Baker N, Munday JC, de Koning HP, Horn D, Maser P. 2015. Chimerization at the AQP2-AQP3 locus is the genetic basis of melarsoprol-pentamidine cross-resistance in clinical *Trypanosoma brucei gambiense* isolates. Int J Parasitol Drugs Drug Resist 5:65–8.

38. Imamura H, Downing T, Van den Broeck F, Sanders MJ, Rijal S, Sundar S, Mannaert A, Vanaerschot M, Berg M, De Muylder G, Dumetz F, Cuypers B, Maes I, Domagalska M, Decuypere S, Rai K, Uranw S, Bhattarai NR, Khanal B, Prajapati VK, Sharma S, Stark O, Schonian G, De Koning HP, Settimo L, Vanhollebeke B, Roy S, Ostyn B, Boelaert M, Maes L, Berriman M, Dujardin JC, Cotton JA. 2016. Evolutionary genomics of epidemic visceral leishmaniasis in the Indian subcontinent. Elife 5.

39. Baker N, de Koning HP, Maser P, Horn D. 2013. Drug resistance in African trypanosomiasis: the melarsoprol and pentamidine story. Trends Parasitol 29:110–8.

40. Alsford S, Kemp LE, Kawahara T, Horn D. 2010. RNA interference, growth and differentiation appear normal in African trypanosomes lacking Tudor staphylococcal nuclease. Mol Biochem Parasitol 174:70–3.

41. Dos Santos SC, Teixeira MC, Dias PJ, Sa-Correia I. 2014. MFS transporters required for multidrug/multixenobiotic (MD/MX) resistance in the model yeast: understanding their physiological function through post-genomic approaches. Front Physiol 5:180.

42. Yan N. 2015. Structural Biology of the Major Facilitator Superfamily Transporters. Annu Rev Biophys 44:257–83.

43. Murungi E, Barlow LD, Venkatesh D, Adung’a VO, Dacks JB, Field MC, Christoffels A. 2014. A comparative analysis of trypanosomatid SNARE proteins. Parasitol Int 63:341–8.

44. Alsford S, Turner DJ, Obado SO, Sanchez-Flores A, Glover L, Berriman M, Hertz-Fowler C, Horn D. 2011. High-throughput phenotyping using parallel sequencing of RNA interference targets in the African trypanosome. Genome Res 21:915–24.

45. Ward DM, Pevsner J, Scullion MA, Vaughn M, Kaplan J. 2000. Syntaxin 7 and VAMP-7 are soluble N-ethylmaleimide-sensitive factor attachment protein receptors required for late endosome-lysosome and homotypic lysosome fusion in alveolar macrophages. Mol Biol Cell 11:2327–33.

46. Fernandez-Prada C, Vincent IM, Brotherton MC, Roberts M, Roy G, Rivas L, Leprohon P, Smith TK, Ouellette M. 2016. Different Mutations in a P-type ATPase Transporter in Leishmania Parasites are Associated with Cross-resistance to Two Leading Drugs by Distinct Mechanisms. PLoS Negl Trop Dis 10:e0005171.

47. Andersen JP, Vestergaard AL, Mikkelsen SA, Mogensen LS, Chalat M, Molday RS. 2016. P4-ATPases as Phospholipid Flippases-Structure, Function, and Enigmas. Front Physiol 7:275.

48. Jeacock L, Baker N, Wiedemar N, Maser P, Horn D. 2017. Aquaglyceroporin-null trypanosomes display glycerol transport defects and respiratory-inhibitor sensitivity. PLoS Pathog 13:e1006307.

49. Uzcategui NL, Figarella K, Bassarak B, Meza NW, Mukhopadhyay R, Ramirez JL, Duszenko M. 2013. Trypanosoma brucei aquaglyceroporins facilitate the uptake of arsenite and antimonite in a pH dependent way. Cell Physiol Biochem 32:880–8.

50. Bassarak B, Uzcategui NL, Schonfeld C, Duszenko M. 2011. Functional characterization of three aquaglyceroporins from Trypanosoma brucei in osmoregulation and glycerol transport. Cell Physiol Biochem 27:411–20.

51. Jhingran A, Chawla B, Saxena S, Barrett MP, Madhubala R. 2009. Paromomycin: uptake and resistance in Leishmania donovani. Mol Biochem Parasitol 164:111–7.

52. Shalev M, Kondo J, Kopelyanskiy D, Jaffe CL, Adir N, Baasov T. 2013. Identification of the molecular attributes required for aminoglycoside activity against Leishmania. Proc Natl Acad Sci U S A 110:13333–8.

53. Engstler M, Pfohl T, Herminghaus S, Boshart M, Wiegertjes G, Heddergott N, Overath P. 2007. Hydrodynamic flow-mediated protein sorting on the cell surface of trypanosomes. Cell 131:505–15.

54. Alsford S, Field MC, Horn D. 2013. Receptor-mediated endocytosis for drug delivery in African trypanosomes: fulfilling Paul Ehrlich’s vision of chemotherapy. Trends Parasitol 29:207–12.

55. Sundar S, Sinha PK, Rai M, Verma DK, Nawin K, Alam S, Chakravarty J, Vaillant M, Verma N, Pandey K, Kumari P, Lal CS, Arora R, Sharma B, Ellis S, Strub-Wourgaft N, Balasegaram M, Olliaro P, Das P, Modabber F. 2011. Comparison of short-course multidrug treatment with standard therapy for visceral leishmaniasis in India: an open-label, non-inferiority, randomised controlled trial. Lancet 377:477–86.

56. Musa A, Khalil E, Hailu A, Olobo J, Balasegaram M, Omollo R, Edwards T, Rashid J, Mbui J, Musa B, Abuzaid AA, Ahmed O, Fadlalla A, El-Hassan A, Mueller M, Mucee G, Njoroge S, Manduku V, Mutuma G, Apadet L, Lodenyo H, Mutea D, Kirigi G, Yifru S, Mengistu G, Hurissa Z, Hailu W, Weldegebreal T, Tafes H, Mekonnen Y, Makonnen E, Ndegwa S, Sagaki P, Kimutai R, Kesusu J, Owiti R, Ellis S, Wasunna M. 2012. Sodium stibogluconate (SSG) & paromomycin combination compared to SSG for visceral leishmaniasis in East Africa: a randomised controlled trial. PLoS Negl Trop Dis 6:e1674.

57. Venkatesh D, Boehm C, Barlow LD, Nankissoor NN, O’Reilly A, Kelly S, Dacks JB, Field MC. 2017. Evolution of the endomembrane systems of trypanosomatids - conservation and specialisation. J Cell Sci 130:1421–1434.

58. Diro E, Blesson S, Edwards T, Ritmeijer K, Fikre H, Admassu H, Kibret A, Ellis SJ, Bardonneau C, Zijlstra EE, Soipei P, Mutinda B, Omollo R, Kimutai R, Omwalo G, Wasunna M, Tadesse F, Alves F, Strub-Wourgaft N, Hailu A, Alexander N, Alvar J. 2019. A randomized trial of AmBisome monotherapy and AmBisome and miltefosine combination to treat visceral leishmaniasis in HIV co-infected patients in Ethiopia. PLoS Negl Trop Dis 13:e0006988.

59. Goyal V, Mahajan R, Pandey K, Singh SN, Singh RS, Strub-Wourgaft N, Alves F, Rabi Das VN, Topno RK, Sharma B, Balasegaram M, Bern C, Hightower A, Rijal S, Ellis S, Sunyoto T, Burza S, Lima N, Das P, Alvar J. 2018. Field safety and effectiveness of new visceral leishmaniasis treatment regimens within public health facilities in Bihar, India. PLoS Negl Trop Dis 12:e0006830.

60. Moreira RA, Mendanha SA, Fernandes KS, Matos GG, Alonso L, Dorta ML, Alonso A. 2014. Miltefosine increases lipid and protein dynamics in Leishmania amazonensis membranes at concentrations similar to those needed for cytotoxicity activity. Antimicrob Agents Chemother 58:3021–8.

61. Currier RB, Cooper A, Burrell-Saward H, MacLeod A, Alsford S. 2018. Decoding the network of *Trypanosoma brucei* proteins that determines sensitivity to apolipoprotein-L1. PLoS Pathog 14:e1006855.

62. Thomson R, Finkelstein A. 2015. Human trypanolytic factor APOL1 forms pH-gated cation-selective channels in planar lipid bilayers: relevance to trypanosome lysis. Proc Natl Acad Sci U S A 112:2894–9.

63. Vanwalleghem G, Fontaine F, Lecordier L, Tebabi P, Klewe K, Nolan DP, Yamaryo-Botte Y, Botte C, Kremer A, Burkard GS, Rassow J, Roditi I, Perez-Morga D, Pays E. 2015. Coupling of lysosomal and mitochondrial membrane permeabilization in trypanolysis by APOL1. Nat Commun 6:8078.

64. Shaw CD, Lonchamp J, Downing T, Imamura H, Freeman TM, Cotton JA, Sanders M, Blackburn G, Dujardin JC, Rijal S, Khanal B, Illingworth CJ, Coombs GH, Carter KC. 2016. In vitro selection of miltefosine resistance in promastigotes of Leishmania donovani from Nepal: genomic and metabolomic characterization. Mol Microbiol 99:1134–48.

65. Coelho AC, Boisvert S, Mukherjee A, Leprohon P, Corbeil J, Ouellette M. 2012. Multiple mutations in heterogeneous miltefosine-resistant *Leishmania major* population as determined by whole genome sequencing. PLoS Negl Trop Dis 6:e1512.

66. Dean S, Sunter JD, Wheeler RJ. 2017. TrypTag.org: A Trypanosome Genome-wide Protein Localisation Resource. Trends Parasitol 33:80–82.

67. Gray KC, Palacios DS, Dailey I, Endo MM, Uno BE, Wilcock BC, Burke MD. 2012. Amphotericin primarily kills yeast by simply binding ergosterol. Proc Natl Acad Sci U S A 109:2234–9.

68. Belenky P, Camacho D, Collins JJ. 2013. Fungicidal drugs induce a common oxidative-damage cellular death pathway. Cell Rep 3:350–8.

69. Purkait B, Kumar A, Nandi N, Sardar AH, Das S, Kumar S, Pandey K, Ravidas V, Kumar M, De T, Singh D, Das P. 2012. Mechanism of amphotericin B resistance in clinical isolates of *Leishmania donovani*. Antimicrob Agents Chemother 56:1031–41.

70. Pountain AW, Weidt SK, Regnault C, Bates PA, Donachie AM, Dickens NJ, Barrett MP. 2019. Genomic instability at the locus of sterol C24-methyltransferase promotes amphotericin B resistance in Leishmania parasites. PLoS Negl Trop Dis 13:e0007052.

71. Alsford S, Kawahara T, Glover L, Horn D. 2005. Tagging a *T. brucei* RRNA locus improves stable transfection efficiency and circumvents inducible expression position effects. Mol Biochem Parasitol 144:142–8.

72. Schumann Burkard G, Jutzi P, Roditi I. 2011. Genome-wide RNAi screens in bloodstream form trypanosomes identify drug transporters. Mol Biochem Parasitol 175:91–4.

73. Alsford S, Horn D. 2008. Single-locus targeting constructs for reliable regulated RNAi and transgene expression in *Trypanosoma brucei*. Mol Biochem Parasitol 161:76–9.

74. Raz B, Iten M, Grether-Buhler Y, Kaminsky R, Brun R. 1997. The Alamar Blue assay to determine drug sensitivity of African trypanosomes (*T. b. rhodesiense* and *T. b. gambiense*) in vitro. Acta Trop 68:139–47.

75. Langmead B, Trapnell C, Pop M, Salzberg SL. 2009. Ultrafast and memory-efficient alignment of short DNA sequences to the human genome. Genome Biol 10:R25.

76. Li H, Handsaker B, Wysoker A, Fennell T, Ruan J, Homer N, Marth G, Abecasis G, Durbin R, Genome Project Data Processing S. 2009. The Sequence Alignment/Map format and SAMtools. Bioinformatics 25:2078–9.

77. Rutherford K, Parkhill J, Crook J, Horsnell T, Rice P, Rajandream MA, Barrell B. 2000. Artemis: sequence visualization and annotation. Bioinformatics 16:944–5.

78. Huson DH, Scornavacca C. 2012. Dendroscope 3: an interactive tool for rooted phylogenetic trees and networks. Syst Biol 61:1061–7.

79. Tsirigos KD, Peters C, Shu N, Kall L, Elofsson A. 2015. The TOPCONS web server for consensus prediction of membrane protein topology and signal peptides. Nucleic Acids Res 43:W401–7.

80. Redmond S, Vadivelu J, Field MC. 2003. RNAit: an automated web-based tool for the selection of RNAi targets in Trypanosoma brucei. Mol Biochem Parasitol 128:115–8.

81. Alsford S, Currier RB, Guerra-Assuncao JA, Clark TG, Horn D. 2014. Cathepsin-L can resist lysis by human serum in *Trypanosoma brucei brucei*. PLoS Pathog 10:e1004130.

