## Supplemental Figures 1-7 for "Chemogenomic profiling of anti-leishmanial efficacy and resistance in the related kinetoplastid parasite *Trypanosoma brucei*"

### Figure S1 (relates to Figure 2 and Table S1)

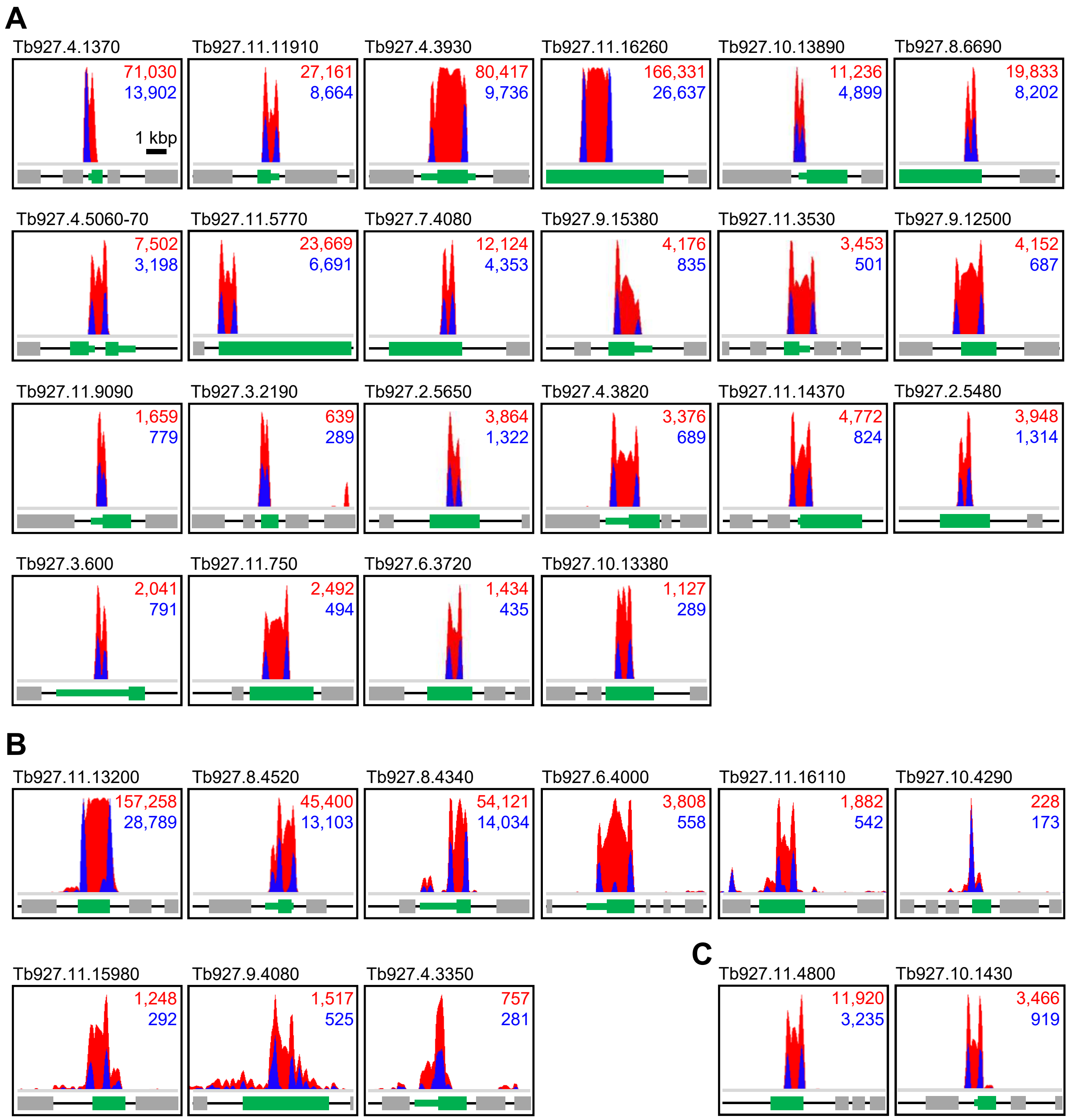


**Candidate anti-leishmanial drug efficacy determinants identified by *T. brucei* RNAi library selection.** Total (red) and RNAi construct-specific 14mer-containing (blue) reads mapping to individual loci following BSF *T. brucei* RNA library selection in paromomycin (A), amphotericin-B (B) and miltefosine (C). Targeted open reading frames highlighted in green; flanking open reading frames coloured grey. Where a substantial number of reads target regions outside the open reading frame, the predicted untranslated region is highlighted by a narrow green bar. See Table S1 for further details.

### Figure S2 (relates to Figure 4)

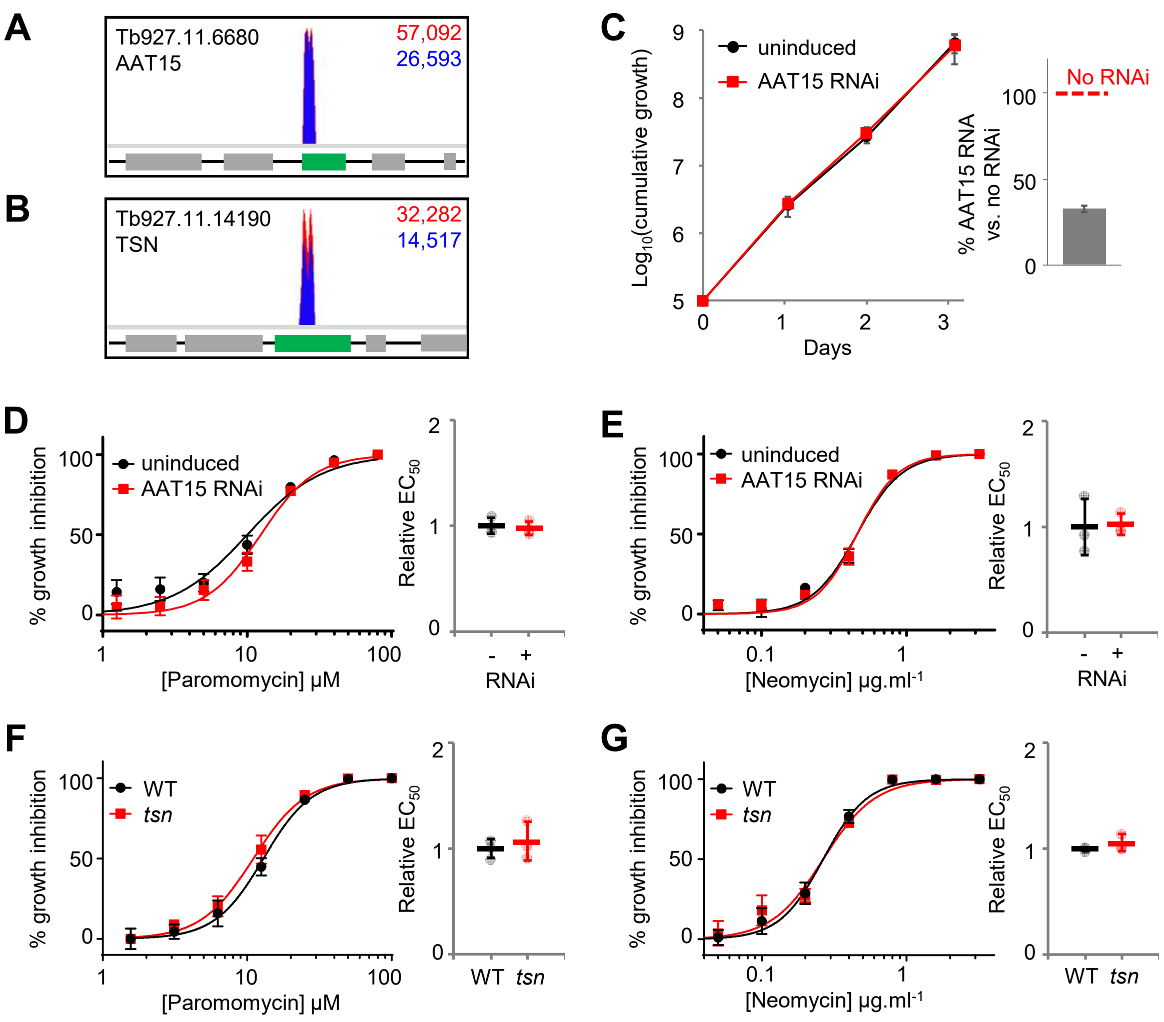


**Neither AAT15 (Tb927.11.6680) depletion nor Tudor Staphylococcal nuclease (Tb927.11.14190) deletion affects aminoglycoside efficacy against BSF *T. brucei* over 72 hours.** A, B) Total (red) and RNAi construct-specific 14mer-containing (blue) reads mapping to *Tb927.11.6680* (A) and *Tb927.11.14190* (B) following paromomycin selection. Targeted open reading frames highlighted in green; flanking open reading frames coloured grey. C) *T. brucei* population growth following AAT15 RNAi knockdown. Inset: RNA depletion was confirmed by RT-qPCR following 24-hour induction in 1 μg.ml$^{-1}$ tetracycline. D, E) Representative paromomycin and neomycin $EC_{50}$ assays following AAT15 RNAi knockdown induced in 1 μg.ml$^{-1}$ tetracycline. F, G) Representative paromomycin (D) and neomycin (E) $EC_{50}$ assays comparing wild type and *Tb927.11.14190* null (*tsn*) BSF *T. brucei*. Inset charts summarise data from three independent biological replicates. Individual growth (C) and $EC_{50}$ (D-G) assays were carried out in triplicate and quadruplicate, respectively. Error bars represent standard deviation.

### Figure S3 (relates to Figure 4)

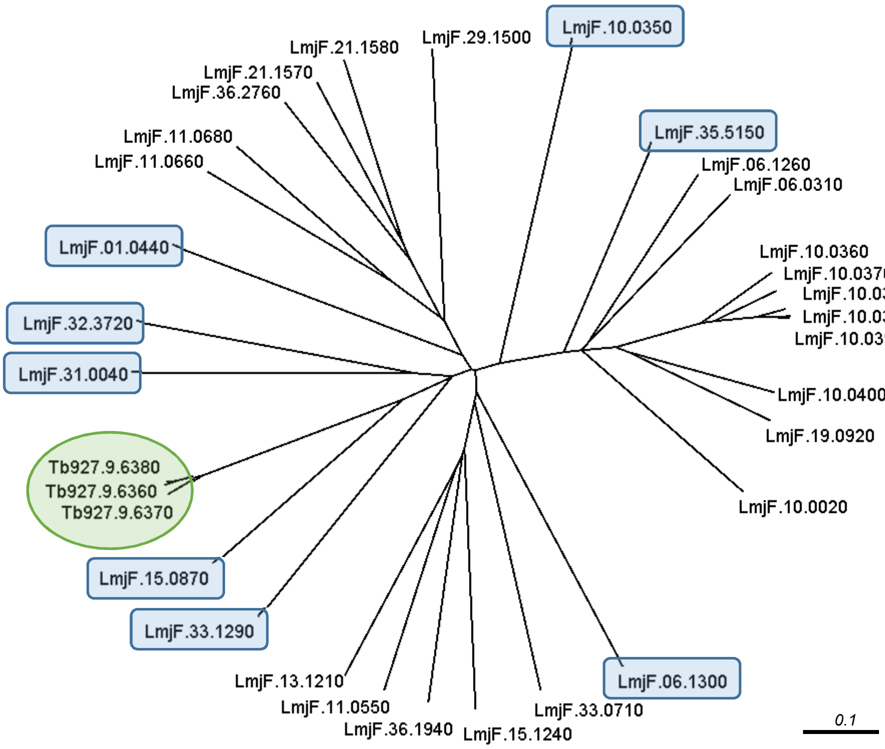


***Tb927.9.6360-80* clusters with the syntenic *LmjF.15.0870*.** Twenty nine open reading frames annotated 'major facilitator' or 'MFS' in the *L. major* Friedlin reference genome were aligned with the Tb927.9.6360-80 open reading frames using Clustal Omega (https://www.ebi.ac.uk/Tools/msa/clustalo/). The unrooted neighbour joining phylogenetic tree was formatted in Dendroscope 3 (http://dendroscope.org/).

### Figure S4 (relates Figure 4)

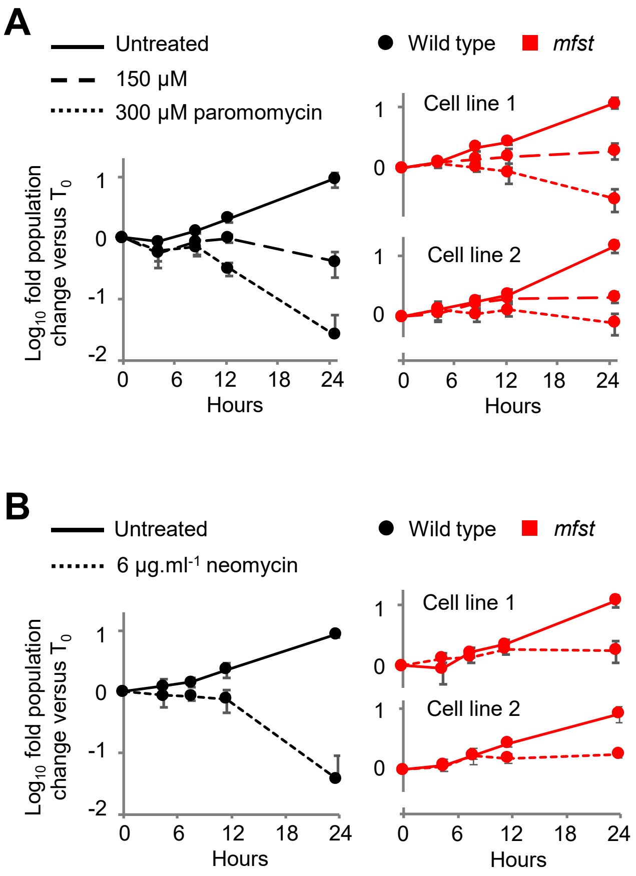


***MFST* locus null *T. brucei* exhibit enhanced tolerance to high concentration aminoglycosides.** Relative population growth of wild type (WT) and *MFST* locus null (*mfst*) *T. brucei* in A) paromomycin and B) neomycin at >$EC_{99}$. Assays were carried out in triplicate. Error bars represent standard deviation.

### Figure S5 (relates to Figure 6)

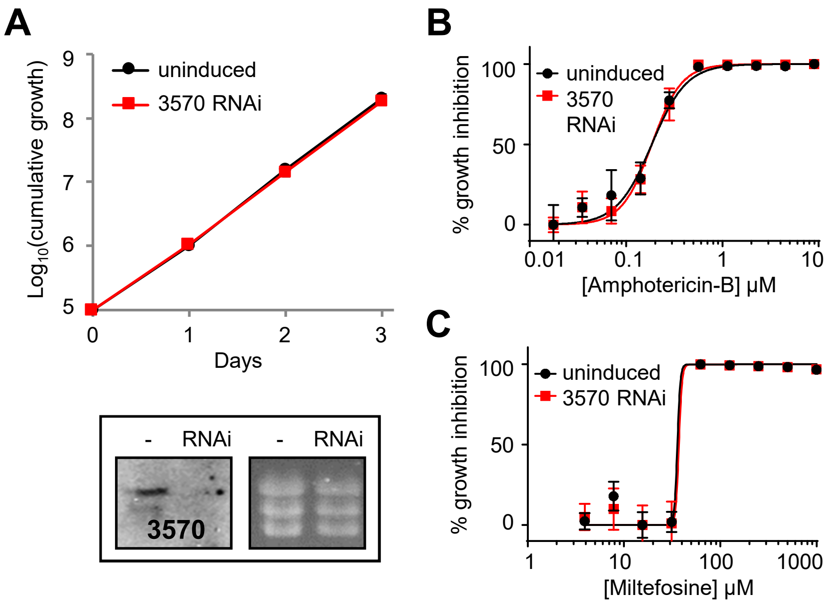


**Tb927.5.3570 does not contribute to the efficacy of amphotericin-B or miltefosine against *T. brucei*.** A) *T. brucei* population growth following Tb927.5.3570 RNAi knockdown. Inset: confirmation of RNAi knockdown by northern blot; ethidium bromide stained gel shown as a loading control. B, C) Representative amphotericin-B and miltefosine $EC_{50}$ assays following Tb927.5.3570 RNAi knockdown. RNAi inductions were carried out in 1 μg.ml$^{-1}$ tetracycline. Individual growth (A) and $EC_{50}$ (B, C) assays were carried out in triplicate and quadruplicate, respectively. Error bars represent standard deviation.

Figure S6 (relates to Figures 7 & 8)

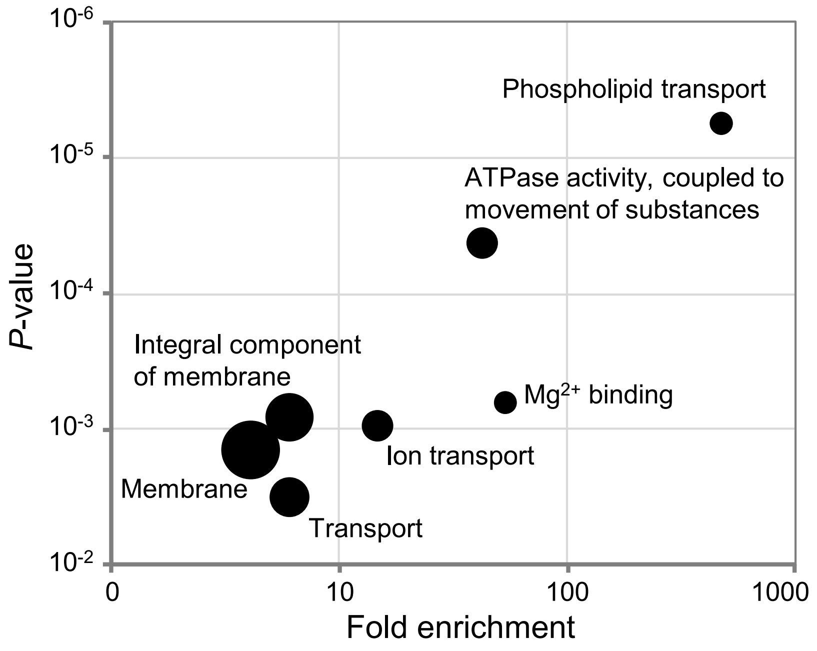


**Gene Ontology analysis of the high confidence hits identified following amphotericin-B RNAi library selection.** Plot generated using the GO analysis tool at TritrypDB.org. Point diameter corresponds to relative number of proteins in each category. See Table S2 for further details.

### Figure S7 (relates to Figures 7 & 8)

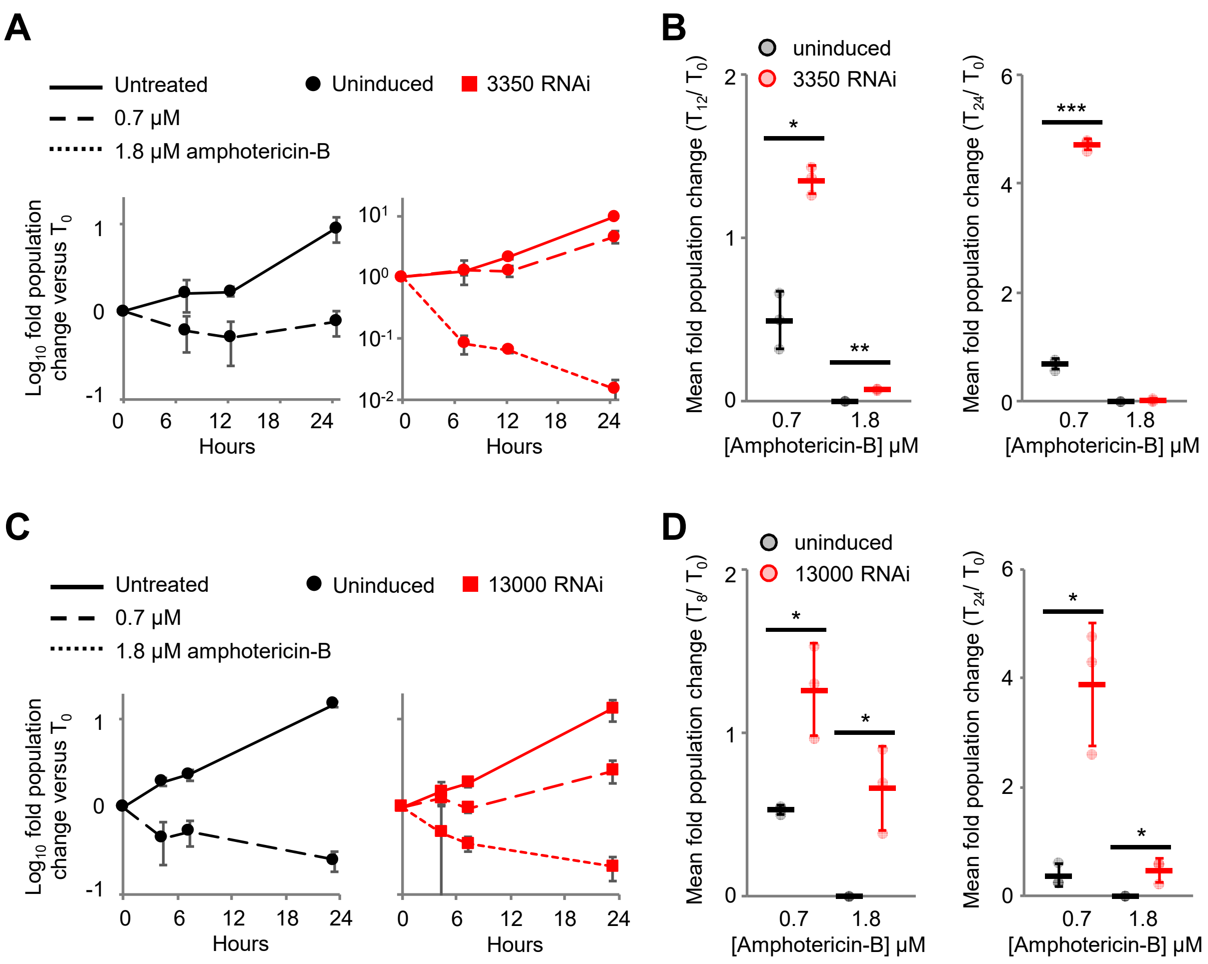


***T. brucei* exhibit enhanced tolerance to high concentration amphotericin-B following flippase depletion.** A, C) Representative assays showing relative population growth in >$EC_{99}$ amphotericin-B following (A) Tb927.11.3350 and (C) Tb927.11.13000 RNAi knockdown. B, D) Relative population growth in >$EC_{99}$ amphotericin-B following (B) Tb927.11.3350 and (D) Tb927.11.13000 RNAi knockdown; data derived from three independent biological replicates. Individual growth assays were carried out in triplicate. Error bars represent standard deviation. *P*-values derived from Student's *t*-test (* <0.05; *** <0.001). RNAi inductions were carried out in 1 μg.ml$^{-1}$ tetracycline.
